# Strong Protection by Bazedoxifene Against Chemically-Induced Ferroptotic Neuronal Death *In Vitro* and *In Vivo*

**DOI:** 10.1101/2024.05.26.595988

**Authors:** Xiangyu Hao, Yifan Wang, Yong Xiao Yang, Lixi Liao, Tongxiang Chen, Pan Wang, Xiaojun Chen, Bao Ting Zhu

## Abstract

Ferroptosis is a form of regulated cell death characterized by excessive iron-dependent lipid peroxidation. Ferroptosis can be induced in cultured cells by exposure to certain chemicals (*e.g.*, erastin and RSL3). Recently it was shown that protein disulfide isomerase (PDI) is a mediator of chemically-induced ferroptosis and also a target for ferroptosis protection. In this study, we find that bazedoxifene (BAZ), a selective estrogen receptor modulator with reported neuroprotective actions in humans, can inhibit PDI function and also strongly protect against chemically-induced ferroptosis in cultured neuronal cells. We find that BAZ can directly bind to PDI *in vitro* and in intact neuronal cells, and also can inhibit PDI’s catalytic activity. Computational modeling analysis reveals that BAZ forms a hydrogen bond with PDI-His256. Inhibition of PDI by BAZ markedly reduces nNOS and iNOS dimerization and NO accumulation, which have recently been shown to play a crucial role in mediating chemically-induced ferroptosis. In addition, the direct antioxidant activity of BAZ may also partially contribute to its protective effect against chemically-induced ferroptosis. Behavioral analysis shows that mice treated with BAZ are strongly protected against kainic acid-induced memory deficits and hippocampal neuronal damage *in vivo*. In conclusion, the results of this study demonstrate that BAZ is an inhibitor of PDI and can strongly prevent chemically-induced ferroptosis in hippocampal neurons both *in vitro* and *in vivo*. These observations offer a novel, estrogen receptor-independent mechanism for the recently-reported neuroprotective actions of BAZ in humans.

**SIGNIFICANCE STATEMENT:** Ferroptosis is an iron- and lipid peroxidation-dependent form of regulated cell death. Recent evidence has shown that protein disulfide isomerase (PDI) is an important mediator of chemically-induced ferroptosis and also a new target for ferroptosis protection. We find that bazedoxifene is an inhibitor of PDI, which can strongly protect against chemically-induced ferroptotic neuronal death *in vitro* and *in vivo*. Additionally, the molecular mechanism of PDI□bazedoxifene binding interaction is defined. This work provides evidence for an estrogen receptor-independent, PDI-mediated mechanism of neuroprotection by bazedoxifene.

## INTRODUCTION

Ferroptosis is a new form of regulated cell death driven by glutathione (GSH) depletion and/or reduction in glutathione peroxidase 4 (GPX4) activity, both of which can lead to accumulation of cellular lipid reactive oxygen species (ROS), and ultimately an iron-dependent oxidative cell death (ferroptosis) (1-7). Erastin is an inhibitor of the amino acid antiporter system Xc^D^, which mediates the influx of extracellular cystine (1,8). Depletion of intracellular cystine leads to reduced cysteine and GSH biosynthesis, and ultimately ferroptotic cell death resulting from GSH depletion (1,8). In comparison, RSL3 is an inhibitor of GPX4, and can effectively induce ferroptosis resulting from accumulation of cellular lipid-ROS (9-11). There is emerging evidence showing that ferroptosis is involved in a number of human diseases, such as neurodegeneration, ischemic or chemically-induced liver injury, and cancer (11,12). Pharmacological manipulation of ferroptosis has been suggested as a potential therapeutic strategy (13).

Protein disulfide isomerase (PDI or PDIA1) is a prototype of the PDI family proteins, which is a ubiquitous dithiol/disulfide oxidoreductase of the thioredoxin superfamily (14-16). PDI is primarily localized in the endoplasmic reticulum of mammalian cells, although a small fraction of this protein is also found in the nucleus, cytosol, mitochondria, plasma membrane and extracellular space (17-19). PDI is involved in protein processing by catalyzing the formation of intra- and inter-molecular disulfide bridges in proteins (17). Earlier we have demonstrated that PDI plays an important role in mediating chemically-induced, GSH depletion-associated oxidative cytotoxicity in HT22 cells, an immortalized mouse hippocampal neuronal cell line (20). Mechanistically, we have shown that glutamate- or erastin-induced GSH depletion can lead to activation of PDI, which then mediates ferroptosis by catalyzing NOS dimerization, followed by accumulation of cellular NO, ROS and lipid-ROS, and ultimately oxidative cell death (ferroptosis) (20,21). Pharmacological inhibition of PDI’s catalytic function or selective PDI knockdown each can effectively abrogate erastin-induced ferroptosis in HT22 cells (21). In addition, our recent study has shown that PDI plays a similar role in mediating erastin-induced, GSH depletion-associated ferroptotic cell death in the estrogen receptor-negative MDA-MB-231 human breast cancer cells (22). These studies jointly reveal that PDI plays an important role in mediating chemically-induced, GSH depletion-associated ferroptosis, which also highlights PDI as a potential cellular target for protection against ferroptotic cell death.

Bazedoxifene (BAZ), a synthetic selective estrogen receptor modulator (SERM) (structure shown in **Fig. 1**), has been approved for use in the U.S. for treatment of osteoporosis in postmenopausal women (23,24). BAZ is presently also being studied for its anticancer activity in humans, including human breast cancer (25). In addition to its anticancer activity as a monotherapy, BAZ has been shown to enhance the chemotherapeutic efficacy of clinically-used anticancer agents such as paclitaxel, cisplatin, oxaliplatin and palbociclib in multiple neoplasms (26-29). Besides, there have been studies in recent years reporting that some of the synthetic SERMs (including BAZ) have a protective effect in neurons and glial cells against toxic insults (30-32). It has been suggested that the mechanism of the protective action of SERMs may involve activation of the estrogen receptors and the G protein-coupled receptor for estrogens (GRP30), through increased expression of antioxidants and activation of kinase-mediated survival signaling pathways (33). However, the exact mechanism by which BAZ exerts its neuroprotective action is still not clear at present, which is the focus of our present study.

**Figure 1.**
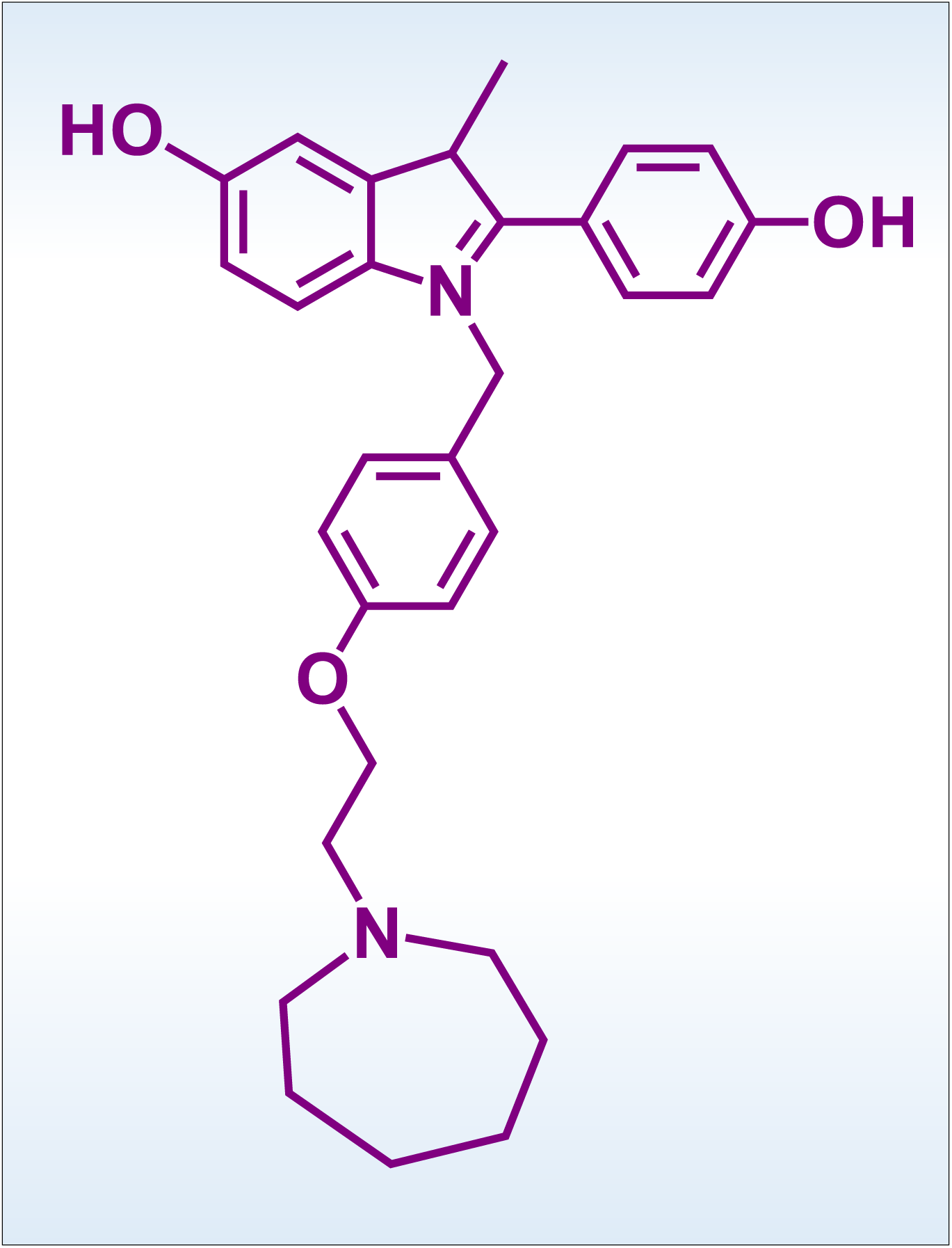
Chemical structure of bazedoxifene (BAZ).

The present study sought to determine the neuroprotective effect of BAZ against chemically-induced ferroptotic neuronal death using both *in vitro* and *in vivo* models, with a focus on determining whether BAZ exerts its neuroprotection through altering PDI function and its downstream signaling pathway. We find that BAZ has a strong protective effect against chemically-induced ferroptosis in mouse neuronal cells both *in vitro* and *in vivo*. Mechanistically, BAZ can effectively inhibit chemically-induced ferroptosis through its binding interaction with PDI and inhibition of PDI’s catalytic activity, which then suppresses NOS dimerization and NO accumulation, followed by reduced accumulation of cellular ROS/lipid-ROS and ultimately ferroptotic cell death. In addition, it is believed that the direct antioxidant activity of BAZ may also contribute to its neuroprotection against chemically-induced ferroptosis.

## Experimental Procedures

### Chemicals

BAZ (#HY-A0031) and kainic acid (#HY-N2309) were obtained from MedChemExpress (Monmouth Junction, NJ, USA), and dissolved in pure dimethyl sulfoxide (DMSO) or saline, respectively, to prepare their stock solutions (at 10 mM). Erastin (#S7242) and RSL3 (#S8155) were purchased from Selleck Chemicals (Houston, TX, USA), and their stock solutions (at 1 mM) were prepared in DMSO. 2’,7’-Dichlorodihydrofluorescein diacetate (DCFH-DA, #S0033S) and 3-amino,4-aminomethyl-2’,7’-difluorescein diacetate (DAF-FM-DA, #S0019S) were purchased from Beyotime Biotechnology (Shanghai, China). MitoSOX (#M36008), MitoTracker green (#M7514) and BODIPY-581/591-C11 (#D3861) were obtained from ThermoFisher (Waltham, MA, USA).

### Cell culture and cell viability assay

The HT22 mouse hippocampal neuronal cells and MDA-MB-231 human breast cancer cells were obtained from the Cell Bank of Chinese Academy of Sciences (Shanghai, China), and were maintained in DMEM supplemented with 10% (*v*/*v*) fetal bovine serum (FBS, ThermoFisher, Waltham, MA, USA) and antibiotics (containing 100IZU/mL penicillin and 100IZμg/mL streptomycin; Sigma-Aldrich). Cells were cultured at 37IZ under 5% CO_2_. Cell viability was determined by the MTT assay as described earlier (22). Cells were authenticated by STR profiling and routinely tested for mycoplasma contamination.

### Staining of live and dead cells

Live and dead cells in culture were distinguished using the calcein-AM and propidium iodide (PI) double staining method, according to the instructions of the manufacturer (Solarbio, Beijing, China). Briefly, after treatment of HT22 cells with selected chemicals, 1 µM calcein-AM and 2.5 μM PI were added to the culture medium and the cells were cultured for an additional 30 min at 37°C in the dark. Images of the cultured cells were then taken using a Nikon Eclipse Ti-U inverted microscope (Nikon, Tokyo, Japan).

### Quantitative real-time polymerase chain reaction (q-PCR)

Total RNAs in cells were sequentially extracted with the TRIzol reagent (#15596018; Invitrogen, Waltham, Massachusetts, USA) and chloroform, precipitated with isopropyl alcohol, and then washed with 75% ethanol to dissolve RNAs in the RNase-free sterile water. The cDNAs were synthesized with the Hifair III 1^st^ Strand cDNA Synthesis Kit (#R312, Vazyme Biotech Co., Ltd, Nanjing, China). Subsequently, q-PCR was performed using PerfectStart Green qPCR SuperMix (#AQ602, TransGen Biotech, Beijing, China) on an Applied Biosystems QuantStudio 3 (ThermoFisher, Waltham, MA, USA). Relative gene expression was calculated using the 2−ΔΔC_t_ method, with GAPDH serving as an internal control. All primers (sequences are shown in **Table 1**) were synthesized by Sangon Biotech (Shanghai, China).

**Table 1.**
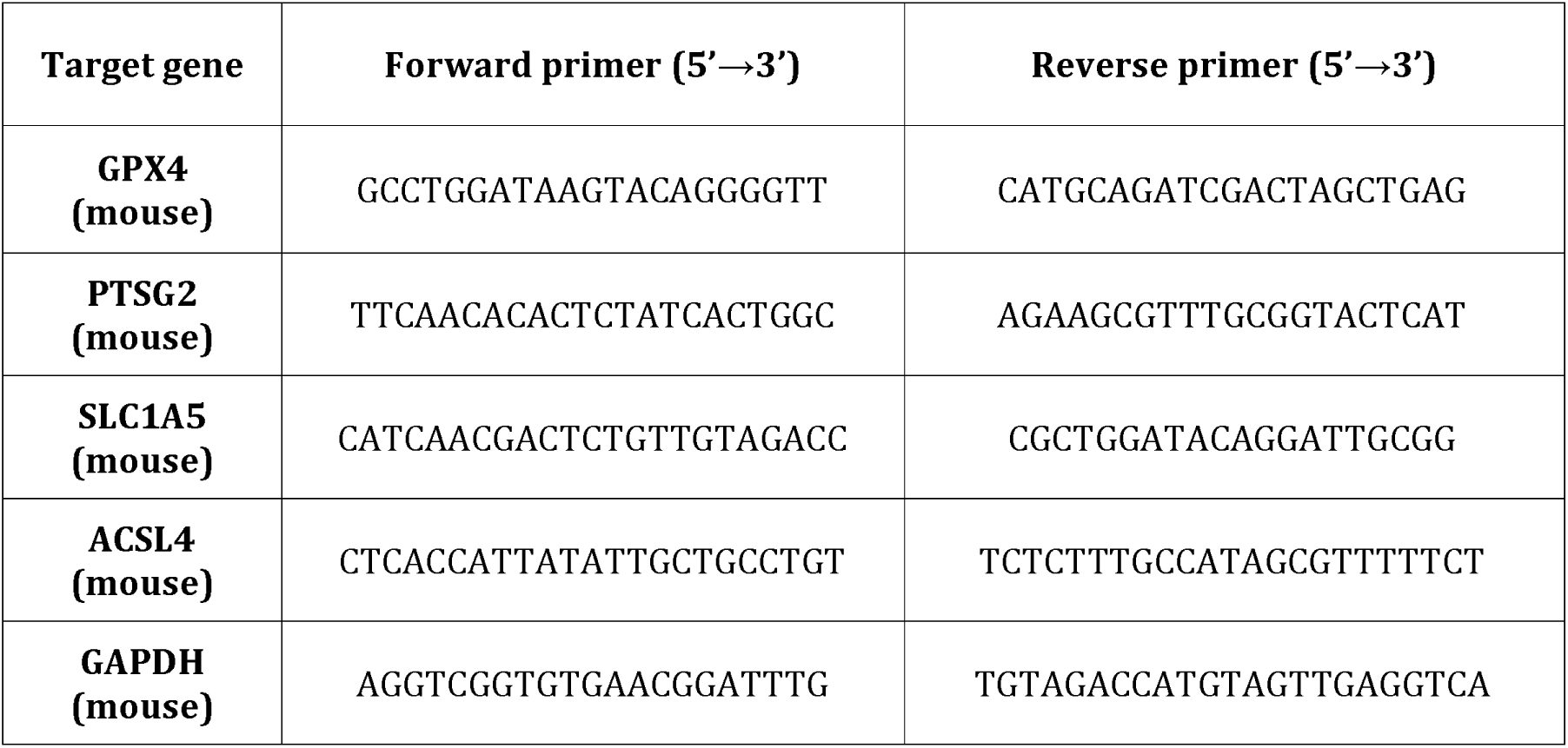
Primer sequences used in the q-PCR analysis.

### Measurement of cellular NO, ROS and mitochondrial ROS by fluorescence microscopy

Cells were plated in 24-well places at a density of 5 IZ 10^4^ per well and treated with drugs or chemicals for selected durations. For fluorescence staining, cells were first washed twice with HBSS and then incubated with DAF-FM-DA (5 μM, for cellular NO), DCFH-DA (5 μM, for cellular ROS), MitoSOX (5 μM, for mitochondrial ROS) or MitoTracker green (5 μM) in 200 μL DMEM (free of phenol red and serum) for 20 min at 37°C. Following three washes with HBSS, fluorescence images were captured using an AXIO fluorescence microscope (Carl Zeiss Corporation, Germany).

### Measurement of cellular NO, ROS and lipid-ROS by flow cytometry

Cells were seeded in 6-well plates at a density of 15 × 10^4^ cells/well 24 h before treatment with different drugs. Following drug treatment, cells were trypsinized, collected and suspended in phosphate-buffered saline (PBS). Cells were then centrifuged, and the resulting cell pellets were resuspended in DMEM (free of phenol red and serum) containing 5 μM of DAF-FM-DA, DCFH-DA or BODIPY-581/591-C11. After a 20-min incubation at 37°C, the cells were washed three times with HBSS to remove any remaining fluorescent dyes. Levels of cellular NO, ROS and lipid-ROS were measured using flow cytometry (Beckman Coulter, Brea, CA, USA) and analyzed using the FlowJo software (FlowJo, LLC, Ashland, USA).

### Measurement of lipid-ROS by confocal microscopy

Cells were seeded at a density of 10 IZ 10^4^ per well on coverslips placed inside the 12-well plates. Twenty-four h later, cells were treated with selected chemicals as indicated. Coverslips were then washed in HBSS and incubated in HBSS containing BODIPY-581/591-C11 (5 μM) for 20 min at 37°C. Coverslips were then mounted on microscope slides for visualization. Cells on the slides were visualized using a LSM 900 confocal laser scanning microscope (LSM 900; Carl Zeiss, Oberkochen, Germany), and images were analyzed with the Zen software (Carl Zeiss).

### Immunoblotting assay

Following drug treatment of HT22 cells as indicated, the cells were lysed on ice for 15 min with the RIPA buffer (#P0013B, Beyotime Biotechnology, Shanghai China) containing a protease inhibitor cocktail (Selleck Chemicals, Houston, TX, USA). After centrifugation for 15 min at 4°C at 13,000 *rpm*, the supernatants were mixed with 5IZ SDS sample buffer (Beyotime Biotechnology, Shanghai, China) and the proteins of interest were separated by electrophoresis with SDS-PAGE. For immunoblot analysis of the dimeric and monomeric forms of nNOS and iNOS, protein samples were prepared with a non-reducing sample buffer without heating (34). The primary antibodies against PDI (#3501S; Cell Signaling Technology, Beverly, MA, USA), nNOS and iNOS (#ab76067 and #ab178945, respectively; Abcam, Cambridge, MA, USA) and the mouse anti-rabbit (#3678S) and the anti-mouse HRP-conjugated secondary antibodies (#7076S) were from Cell Signaling Technology (Beverly, MA, USA).

### Cellular thermal shift assay (CETSA)

CETSA was conducted according to the protocols described earlier (35,36). Briefly, after the cells were treated with the vehicle or indicated chemicals for 3IZh, they were washed with ice-cold PBS, harvested by trypsinization, centrifuged and re-suspended in PBS supplemented with a protease inhibitor cocktail (Selleck Chemicals, Houston, TX, USA). Equal amounts of cell suspensions were aliquoted into 0.2IZmL PCR microtubes.

Subsequently, aliquots of the cell suspension were heated in a Ristretto Thermal Cycler (VWR, Darmstadt, Germany) at the indicated temperatures for 3IZmin, followed by cooling for 3IZmin at room temperature. Finally, the cells were lysed using three cycles of freeze–thawing, and the soluble fractions were isolated by centrifugation and analyzed by SDS-PAGE followed by Western blotting as described above. For isothermal-dose-response-CETSA (ITDR_CETSA_), the fold of change in PDI (which was normalized to the β-actin control) was plotted as a function of temperature to generate the PDI melting curves for selected treatments.

### Protein expression and purification

The mutant PDI-Ala256 protein was prepared from the full-length wild-type human PDI cDNA using the QuikChange II XL Site-Directed Mutagenesis Kit (Agilent Technologies). For protein purification, PCR products of the wild-type PDI-His256 and the mutant PDI-Ala256 were subcloned into pET28a, and proteins were expressed in the *E. coli* strain JM109 (DE3) cultured in Luria-Bertani (LB) medium. After the bacteria were cultured at 37°C to reach an OD value of approximately 0.8, then 1 mM isopropyl-β-*D*-1-thiogalactopyranoside (IPTG) was added into LB medium, and the bacteria were further cultured at 22°C for 8 h. The bacteria were harvested by centrifugation at 5000 × g for 30 min at 4°C, and lysed in the presence of a protease inhibitor cocktail. The supernatant was isolated by centrifugation at 20,000 × g for 30 min and then incubated with Ni-NTA agarose (QIAGEN) at 4°C for 1 h. The columns were washed and eluted with 20 mM imidazole. The proteins were concentrated and analyzed by SDS-PAGE.

### Assay of PDI catalytic activity and its inhibition by BAZ

The effect of BAZ on the catalytic activity of purified wild-type PDI-His256 and mutant PDI-Ala256 was determined by analyzing PDI-mediated aggregation of the insulin B chain as described earlier with some modifications (37,38). Briefly, insulin (125 μM) was incubated in a 96-well plate in 10 mM sodium phosphate buffer (pH 7.4) and 5 mM DTT with or without the recombinant wild-type PDI-His256 or mutant PDI-Ala256 (at 3.5 μM). The aggregation was monitored at 37°C using a Synergy Plate Reader (Biotek, Winooski, VT, USA), with the wavelength set at 650 nm.

### Molecular docking analysis

The PDI□ligand interaction was analyzed using the molecular docking method. The structures of human PDI (PDB code **6I7S**; chain A) (39) and BAZ (extracted from the estrogen receptor□BAZ complex; PDB code **6PSJ**, ligand ID **29S**) (40) were downloaded from the Protein Data Bank (https://www.rcsb.org/) (41) and adopted as the receptor and ligand, respectively.

The structures were processed using the Protein Preparation Wizard in Schrodinger Suite (Maestro 12.8, 2021; Schrodinger LLC, New York, NY, USA). The hydrogen atoms were added, and the protein structures were optimized using OPLS4 force field (42). The protein□ligand docking decoys were generated using Glide-XP (extra precision) in Schrodinger Glide software (43). The nearby torsional minima of the lowest energy binding poses were sampled using Monte Carlo (MC) procedure. The Cα atom of His256 in the ***b’*** domain of PDI was used as the center of the docking grid box with dimensions set at 30 × 30 × 30 Å^3^.

Lastly, three representative scoring functions, X-Score (44), PRODIGY-LIG (45) and Δ_vina_RF_20_ (46) were employed for further filtering of the docking results. The two linear empirical scoring functions, *i.e.*, X-Score (44) and PRODIGY-LIG (45), were developed for calculating protein□ligand binding affinity. The former employs energy and geometric terms such as *van der Waals* energy, hydrogen bonding energy, deformation penalty and hydrophobic effect (44), and the latter uses the number of atomic contacts and electrostatic energy (45). Δ_vina_RF_20_ is a random forest-based method for binding affinity prediction based on 20 descriptors (46).

To investigate the importance of His256 in the interactions between PDI and BAZ, His256 was mutated to alanine (Ala) in the representative predicted structures of the PDI□BAZ complex. The binding energies were predicted using X-Score (44), PRODIGY-LIG (45) and Δ_vina_RF_20_ (46).

### Molecular dynamics simulations

The stability of the binding poses of BAZ in the predicted structures of PDI□BAZ complex was investigated using molecular dynamics (MD) simulations. The complexes were processed using CHARMM-GUI (https://charmm-gui.org/) to generate the topology files (47). The force field parameters of BAZ were generated based on CHARMM general force field (48), those of protein were generated based on CHARMM36m force field (49). The systems were embedded into a rectangular water box extending the solvent 10 Å in ***x***, ***y***, ***z*** directions, and the TIP3P water model (50) was used. K^+^ and Cl^-^ ions with parameters approximated by Roux *et al*. (51) were added to neutralize the charges of the systems. The energy minimization (10000 steps), equilibrium simulation (0.25 ns) in NVT ensemble and production simulation (100 ns) in NPT ensemble were carried out using NAMD (52). The time step and temperature were set to 2 fs and maintained at 300 K using Langevin dynamics (53), respectively.

The periodic boundary conditions were employed, the short-range electrostatic and *van der Waals* interactions were truncated smoothly with a cutoff (12 Å) and a switching function was employed at 10 Å. Long-range electrostatic interaction was estimated by the particle mesh Ewald algorithm (54,55). The pressure in NPT ensemble was maintained at 1 atm by the Langevin piston method (56). The number of hydrogen bonds formed between BAZ and PDI and the contact number and binding energies between BAZ and PDI in the representative conformations along the MD trajectories were calculated to confirm the importance of His256 and the stability of the predicted binding poses.

### *In vivo* animal experiments and drug treatments

The procedures involving the use of live animals described in this study were approved by the Institutional Animal Care and Use Committee (IACUC) of The Chinese University of Hong Kong (Shenzhen), and the guidelines for humane care of animals set forth by the U.S. National Institutes of Health were followed. Male mice (6IZ8 weeks of age), weighing 20IZ30 g, were purchased from Guangdong Charles River Laboratories (Beijing, China). After arrival, the animals were allowed to acclimatize to the new environment for one week before they were used in experimentation. Mice were randomly divided into different experimental groups (n = 6–8) with comparable average body weights.

Kainic acid (3 µL of 0.2 mM solution in saline) was injected into the left and right lateral ventricles using a microliter syringe under anesthesia with Zoletil50 and xylazine (50 and 5 mg/kg, *i.p.*), and the control mice were injected with 3 µL of vehicle (saline) without kainic acid [64]. The bilateral intracerebroventricular (*i.c.v.*) injection parameters were: anterior/posterior, IZ0.5; rostral, ±1.1; and dorsal/ventral, 2.7. The animals were also given *i.p.* injection of a solution (100 μL) containing 0.375, 0.75 or 1.25 mg/mL of BAZ (the solvent is 10% DMSO + 90% corn oil) once every 2 days. The three BAZ doses approximately equal to 1.5, 3 or 5 mg/kg of body weight. The first dose of BAZ was given 24 h before *i.c.v.* injection of kainic acid, and the treatment lasted for 11 days. Note that in this study, the control animals were sham-operated (receiving *i.c.v.* injection of 3 μL saline solution) and also received *i.p.* injections of 100 μL vehicle (containing 10% DMSO and corn oil). For kainic acid alone group, the animals received *i.p.* injections of 100 μL vehicle (10% DMSO and corn oil); for the BAZ alone group, the animals were sham-operated (receiving *i.c.v.* injection of 3 μL saline solution).

### Memory and learning ability tests

The first test used in this study was the classical Y-maze-based method which was intended to determine the degree of memory impairments in mice following different treatments (57,58). The test started 6 days after *i.c.v.* injection of kainic acid. The Y-maze (made of polyvinyl plastic) was a three-arm maze with equal angles between the three arms (30 cm in length, 5 cm in width, and 15 cm in wall height). As depicted in **Fig. 11B**, the test animals were initially placed at the end of one arm, and the sequence and number of arm entries were recorded manually. Based on earlier studies (57,58), the percentage of trials with all three arms represented, *i.e.*, ABC, CAB or BCA (but not ABA, BAB or CBC), was recorded as an alternation for estimation of the short-term memory. Arms were cleaned between tests with 70% ethanol-containing paper towels to remove odors and residues. The alternation score (%) for each mouse is defined as the ratio of the actual number of alternations to the possible number (defined as the total number of arm entries minus two) multiplied by 100 as in the following equation (59):

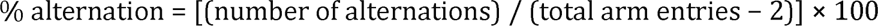

In this study, a new Y-maze-based test was also developed in our laboratory to further evaluate the learning ability and the degree of memory impairments of the same animals. In this method, all tests started from day 10 post *i.c.v.* injection of kainic acid (see **Fig. 11A**), and all mice, including the control mice, were food restricted starting 24 h before the test, receiving on average 2IZ3 g pellet food only. As depicted in **Fig. 11B**, in one of the arms the animal food was placed as a bait. The animal was placed in the starting arm, and then it was allowed to freely explore the Y-maze to find the food (which was placed at the end of another arm) and ate it. Then, the animal was placed back at the same starting place and the observation was repeated again. If the animal went straight to where the food was originally placed and ate the food, it was considered that the animal remembered correctly where the food was. In order to evaluate how well the learning ability and memory of each animal, the number of wrong entries the animal made during the initial 20 trials (10 trials per day, in two consecutive days) and the amount of time the animal took to find and start to eat the food in every trial were recorded. To minimize the random variations in each individual trial, the first three consecutive trials (*i.e.*, trials 1IZ3) were combined to calculate the average value as the first recording, the next three consecutive trials (*i.e.*, trails 4IZ6) were combined to calculate the average value for the second recoding, and the same method was applied to calculate other recording values. The last two trials (*i.e.*, trials 19 and 20) were combined to calculate the average value for the last recoding.

### Immuno- and histochemical staining of the brain sections

To perform immuno- and histochemical staining of the brain sections to determine the degree of brain damage in different treatment groups, separate groups of animals were sacrificed on day 6 post *i.c.v.* injection of kainic acid. Prior to collecting the brain tissues from the animals for analysis, the animals received ketamine and xylazine (50 and 5 mg/kg, *i.p.*) for anesthesia, and then they were perfused with physiological saline (0.9% NaCl) and 4% paraformaldehyde via the abdominal aorta. The collected brain tissues were fixed overnight in 4% paraformaldehyde. After cryoprotection in 30% sucrose/phosphate buffer, the whole brain tissues were frozen in liquid nitrogen and sectioned serially (in 30-µm thickness). Brain sections were collected in 0.1 M neutral phosphate buffer, mounted on slides, then air-dried on a slide warmer at 50°C for at least 0.5 h, and stained with hematoxylin and eosin (H/E) for histological analysis. Three brain hippocampal regions (CA1, CA3 and the dentate gyrus DG) were examined bilaterally as described earlier (60).

The apoptotic DNA degradation in brain tissue slides was determined using the terminal deoxynucleotidyl transferase (TdT)-mediated dUDP-biotin nick end labeling (TUNEL) method according to supplier’s instructions (Biosharp, China). The Fluoro-Jade B staining was performed according to supplier’s instructions (Biosensis, Australia). Briefly, the slides were transferred to a solution of 0.06% potassium permanganate for 10 min on a shaker. The Fluoro-Jade B staining solution was prepared from the 0.1- mg/mL stock solution (in distilled water). After 10 min in the staining solution, the slides were rinsed and placed on a slide warmer until they were dry. Images were captured with a light microscope (Carl Zeiss Corporation, Germany) and cell counting was performed using the ImageJ software.

### Statistical analysis

Most of the experiments described in this study were repeated three times or more to confirm the observations. Data presented in this study are the mean ± S.D. from multiple replicate measurements taken from one representative experiment. Statistical analyses were performed using the GraphPad Prism 7.0 software (GraphPad Software, La Jolla, CA) by using one-way ANOVA coupled with follow-up tests for multiple comparisons. Statistical significance was denoted by *P* < 0.05 (* or ^#^) and *P* < 0.01 (** or ^##^) for significant and very significant differences, respectively. In most cases, * and ** denote the comparison for statistical significance between the control group (cells treated with the vehicle only) and the cells treated with a cell death inducer (such as erastin or RSL3), whereas ^#^ and ^##^ denote the comparison between the cells treated with the cell death inducer alone and the cells jointly treated with the cell death inducer plus a modulating compound (such as BAZ).

## RESULTS

### BAZ prevents erastin-induced ferroptosis and NO/ROS accumulation

#### Protection against ferrotosis

Based on changes in gross morphology and cell viability (MTT assay), treatment of HT22 mouse hippocampal neuronal cells with erastin readily induced cell death in a dose-dependent manner (**Fig. 2A**). Joint treatment of these cells with BAZ (at concentrations ranging from 125 to 2000 nM) abrogated erastin-induced cell death in a concentration-dependent manner (**Fig. 2B**). The protective efficacy of BAZ was very high as 100% protection was readily observed when 500 nM BAZ was present (**Fig. 2B**). Further analysis showed that BAZ effectively attenuated cell death when the live and dead cells were analyzed using calcein AM/PI double staining (**Fig. 2C**).

**Figure 2.**
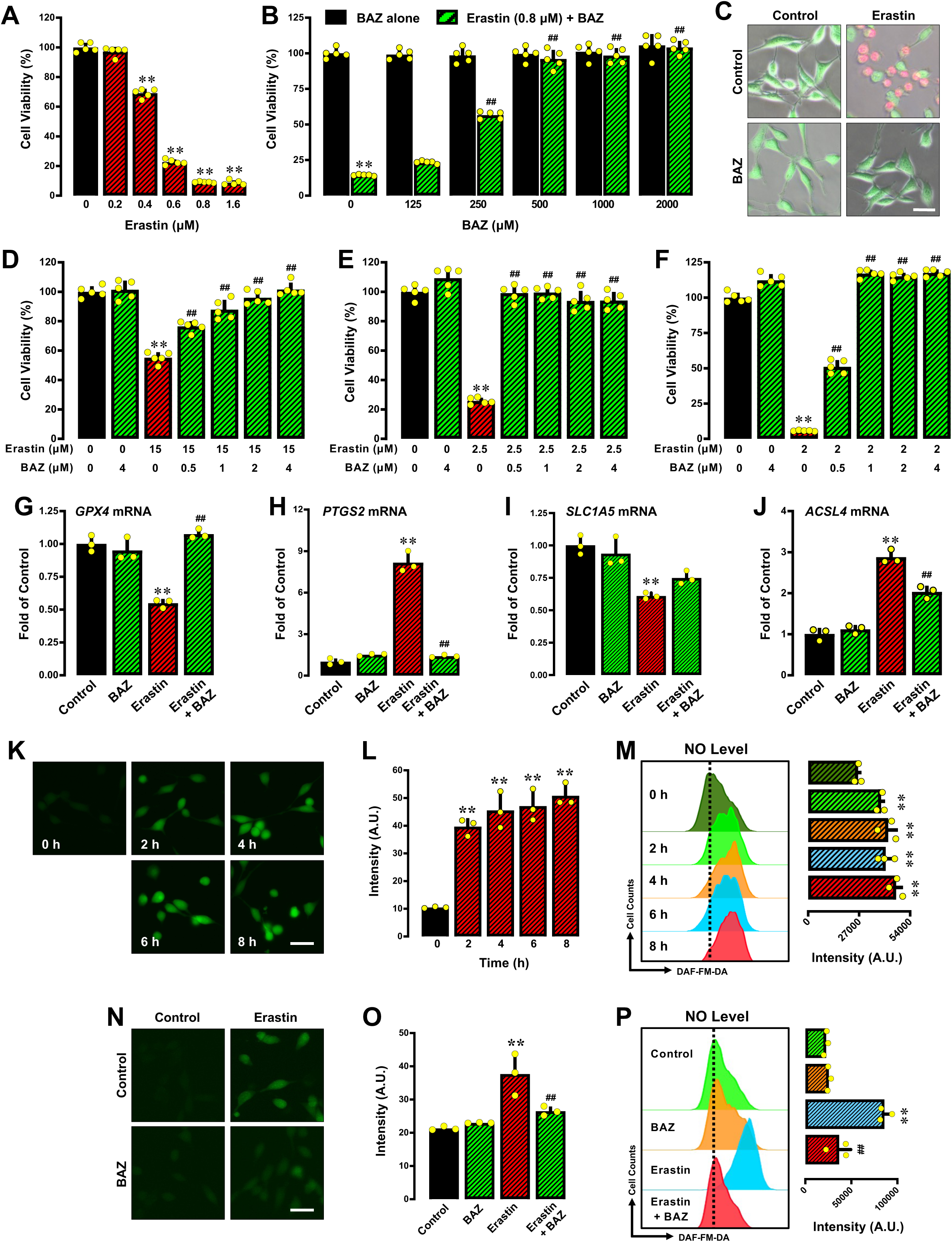
BAZ protects against erastin-induced ferroptosis. **A, B.** HT22 cells were treated with erastin ± BAZ at the indicated concentrations for 24 h. Cell viability was determined by MTT assay and represented as a percentage of control cell viability (n = 5). **C.** Calcein-AM/PI double staining to determine live and dead cells following 8-h treatment of HT22 cells with 0.8 μM erastin ± 1 μM BAZ (fluorescence microscopy images, scale bar = 60 μm). D17F. The MDA-MB-231, H9C2 and BRL-3A cells were treated with erastin ± BAZ at the indicated concentrations for 24 h respectively. Cell viability was determined by MTT assay and represented as a percentage of control cell viability (n = 5). G17J. Changes in mRNA levels of GPX4 (**G**), PTGS2 (**H**), SLC1A5 (**I**) and ACSL4 (**J**) (determined by q-PCR; n = 3) after treatment of HT22 cells with 0.8 μM erastin ± 1 μM BAZ. K17M. Time-dependent induction of cellular NO accumulation following treatment of HT22 cells with 0.8 μM erastin (**K** for fluorescence microscopy images, scale bar = 60 μm; the left panel of **M** for analytical flow cytometry). **L** and the right panel of **M** are the respective quantitative intensity values (n = 3). K17M. Cellular levels of NO after 8-h treatment of HT22 cells with 0.8 μM erastin ± 1 μM BAZ (**K** for fluorescence microscopy images, scale bar = 60 μm; the left panel of **M** for analytical flow cytometry). **L** and the right panel of **M** are the respective quantitative intensity values (n = 3). All quantitative data are presented as mean ± S.D. (* or # *P* < 0.05; ** or ^##^ *P* < 0.01).

We have also determined, for comparison, the protective effect of BAZ against erastin-induced cell death in three additional cell lines, *i.e.*, the estrogen receptor-negative MDA-MB-231 human breast cancer cells, the BRL-3A rat liver cells and the H9C2 rat myocardium cells. We found that BAZ exerted a complete cytoprotection against erastin-induced cell death in these cells (Fig. 2D⍰2F).

In this study, q-PCR analysis of the mRNA levels of representative key ferroptosis markers was performed in HT22 cells treated with erastin ± BAZ, and it was observed that while erastin exposure reduced the mRNA levels of GPX4 (**Fig. 2G**) and SLC1A5 (**Fig. 2I**), it increased the mRNA levels of PTGS2 (**Fig. 2H**) and ACSL4 (**Fig. 2J**). These changes are characteristic for erastin-induced ferroptotic cell death. Joint treatment of cells with BAZ abrogated erastin-induced changes in three of the ferroptosis markers (*GPX4*, *PTGS2* and *ACSL4*) (**Fig. 2G, 2H, 2J**), whereas the change in SLC1A5 was not modest (**Fig. 2I**).

#### Effect on NO and ROS accumulation

Our recent studies showed that erastin-induced cell death in HT22 cells is associated with sequential accumulation of cellular NO, ROS and lipid-ROS (21). By using fluorescence microscopy and flow cytometry analyses, we confirmed that erastin caused accumulation of cellular NO (DAF-FM-DA as a probe) in a time-dependent manner in HT22 cells (Fig. 2K⍰2M), and NO accumulation in erastin-treated cells was abrogated by joint treatment of these cells with 1 µM BAZ (Fig. 2N⍰2P).

Erastin also caused accumulation of cellular ROS (DCFH-DA as a probe) and lipid-ROS (C11-BODIPY as a probe) in a time-dependent manner (**Fig. 3A, 3B, 3F**), and their accumulation was effectively abrogated by BAZ (Fig. 3C⍰3E; 3G⍰3H). In addition, erastin-induced accumulation of mitochondrial ROS (MitoSOX as a probe) was also similarly reduced by treatment with BAZ (**Fig. 3I**).

**Figure 3.**
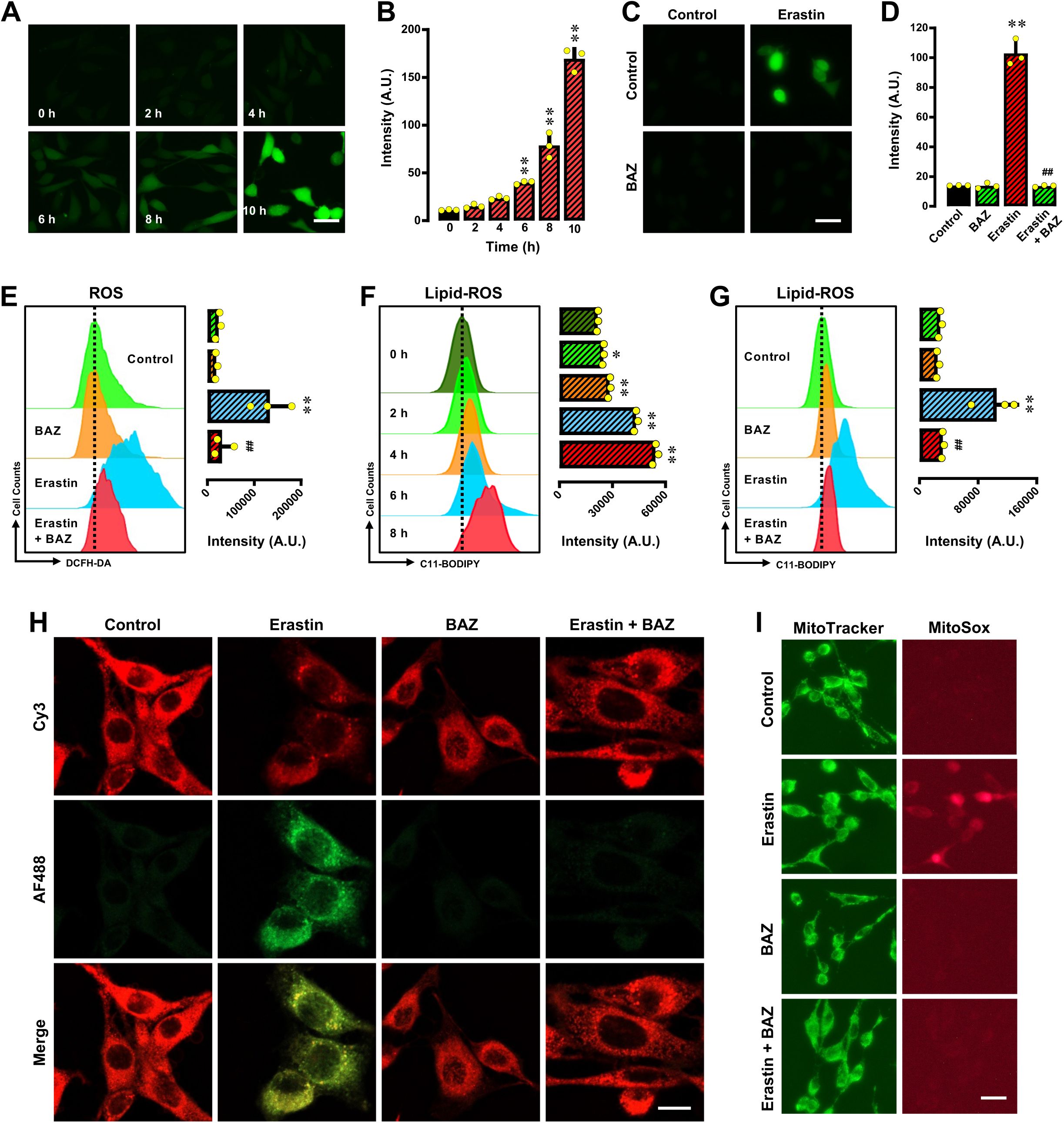
BAZ protects against erastin-induced ferroptosis in HT22 cells. **A, B.** Time-dependent induction of cellular ROS accumulation following 0.8 μM erastin treatment (**A** for fluorescence microscopy images, scale bar = 60 μm, **B** for quantitative intensity values, n = 3). C17E. Cellular levels of ROS after 8-h treatment with 0.8 μM erastin ± 1 μM BAZ (**C** for fluorescence microscopy images, scale bar = 60 μm; the left panel of **E** for analytical flow cytometry). **D** and the right panel of **E** are the respective quantitative intensity values (n = 3). **F.** Time-dependent induction of lipid-ROS accumulation following treatment with 0.8 μM erastin (the left panel for flow cytometry assay; the right panel for quantitative intensity values, n = 3). **G, H.** Cellular levels of lipid-ROS after 8-h treatment with 0.8 μM erastin ± 1 μM BAZ (the left penal of **G** for flow cytometry assay, and **H** for confocal microscopy data, scale bar = 20 μm). The right panel of **G** for respective quantitative intensity values (n = 3). **I.** Levels of mitochondrial ROS after 8-h treatment with 0.8 μM erastin ± 1 μM BAZ (fluorescence microscopy images, scale bar = 60 μm). All quantitative data are presented as mean ± S.D. (* or ^#^ *P* < 0.05; ** or ^##^ *P* < 0.01).

In summary, these results indicate that BAZ has a strong protective effect against erastin-induced ferroptosis in HT22 neuronal cells through attenuation of the accumulation of cellular NO, ROS, lipid-ROS and mitochondrial ROS.

### BAZ protects against RSL3-induced ferroptosis and NO/ROS accumulation

#### Protection against ferrotosis

We have also determined in this study the protective effect of BAZ against RSL3-induced ferroptotic cell death. While treatment of cells with RSL3 alone elicited a dose-dependent loss of cell viability (**Fig. 4A**), a strong protection against RSL3-induced cell death was observed when the cells were jointly treated with BAZ (**Fig. 4B**), and 100% protection was seen when 125 nM BAZ was present. It is evident that the protective effect of BAZ against RSL3-induced cell death has a higher potency than its protection against erastin-induced ferroptosis. Additional experiments using calcein AM/PI double staining confirmed that BAZ effectively attenuated RSL3-induced cell death (**Fig. 4C**).

**Figure 4.**
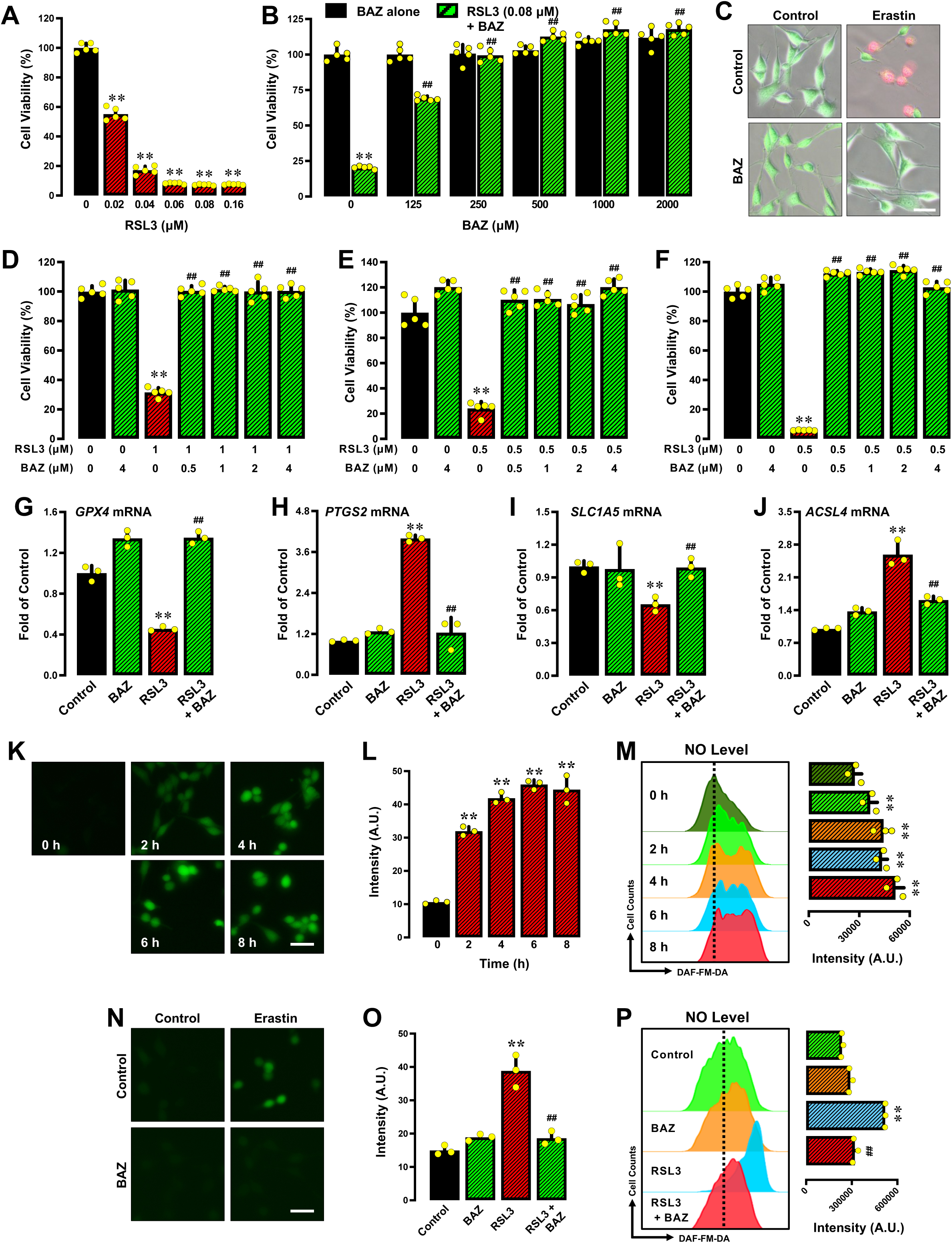
BAZ protects against RSL3-induced ferroptosis. **A, B.** HT22 cells were treated with RSL3 ± BAZ at the indicated concentrations for 24 h. Cell viability was determined by MTT assay and represented as a percentage of control cell viability (n = 5). **C.** Calcein-AM/PI double staining to determine live and dead cells following 8-h treatment of HT22 cells with 0.08 μM RSL3 ± 1 μM BAZ (fluorescence microscopy images, scale bar = 60 μm). D17F. The MDA-MB-231, H9C2 and BRL-3A cells were treated with RSL3 ± BAZ at the indicated concentrations for 24 h respectively. Cell viability was determined by MTT assay and represented as a percentage of control cell viability (n = 5). G17J. Changes in mRNA levels of GPX4 (**G**), PTGS2 (**H**), SLC1A5 (**I**) and ACSL4 (**J**) (determined by q-PCR; n = 3) after treatment of HT22 cells with 0.08 μM RSL3 ± 1 μM BAZ. K17M. Time-dependent induction of cellular NO accumulation following treatment of HT22 cells with 0.08 μM RSL3 (**K** for fluorescence microscopy images, scale bar = 60 μm; the left panel of **M** for analytical flow cytometry). **L** and the right panel of **M** are the respective quantitative intensity values (n = 3). K17M. Cellular levels of NO after 8-h treatment of HT22 cells with 0.08 μM RSL3 ± 1 μM BAZ (**K** for fluorescence microscopy images, scale bar = 60 μm; the left panel of **M** for analytical flow cytometry). **L** and the right panel of **M** are the respective quantitative intensity values (n = 3). All quantitative data are presented as mean ± S.D. (* or # *P* < 0.05; ** or ^##^ *P* < 0.01).

Similar to the observations made with erastin, a strong protection against RSL3-induced cell death was observed when the MDA-MB-231 cells, H9C2 cells and BRL-3A cells were jointly treated with BAZ (Fig. 4D⍰4F), In addition, it was observed that RSL3 decreased the mRNA levels for GPX4 (**Fig. 4G**) and SLC1A5 (**Fig. 4I**) but increased the mRNA levels of PTGS2 (**Fig. 4H**) and ACSL4 (**Fig. 4J**) in HT22 cells. Joint treatment with BAZ abrogated RSL3-induced changes in these mRNA levels (Fig. 4G174J).

#### Effect on NO and ROS accumulation

We also determined the effect of BAZ on RSL3-induced accumulation of cellular NO, ROS/lipid-ROS and mitochondrial ROS. We found that RSL3 caused accumulation of cellular NO in a time-dependent manner (**Fig. 4K⍰4M**), and the presence of 1 µM BAZ effectively abrogated RSL3-induced NO accumulation (Fig. 4N⍰4P). Treatment of HT22 cells with RSL3 also caused accumulation of cellular ROS and lipid-ROS in a time-dependent manner (**Fig. 5A, 5B, 5F**). RSL3-induced cellular ROS and lipid-ROS accumulation was similarly abrogated by BAZ (Fig. 5C⍰5E; 5G⍰5H). In addition, RSL3-induced accumulation of mitochondrial ROS was also abrogated by BAZ (**Fig. 5I**).

**Figure 5.**
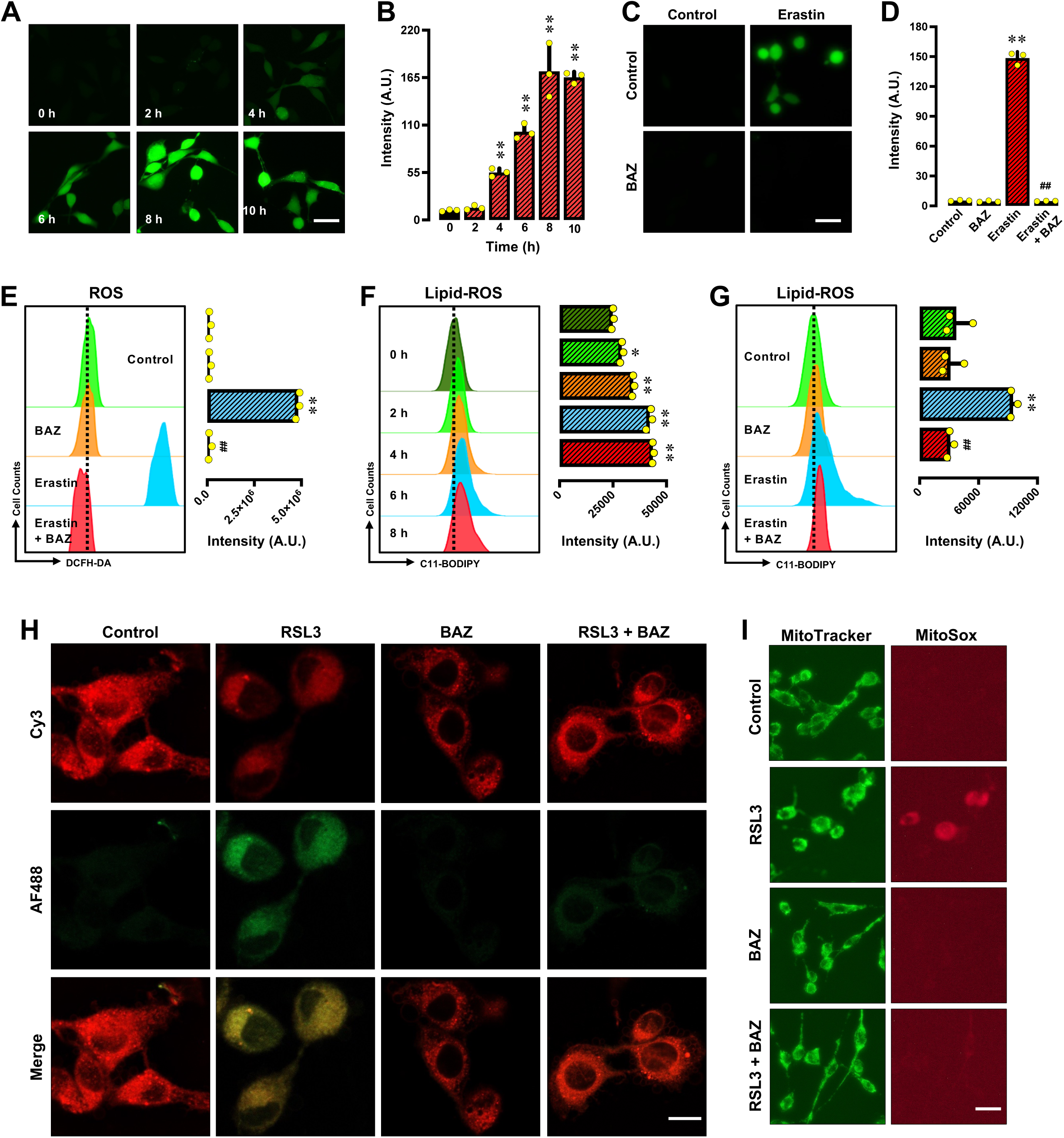
BAZ protects against RSL3-induced ferroptosis in HT22 cells. **A, B.** Time-dependent induction of cellular ROS accumulation following treatment with 0.08 μM RSL3 (**A** for fluorescence microscopy images, scale bar = 60 μm, **B** for quantitative intensity values, n = 3). C17E. Cellular levels of ROS after 8-h treatment with 0.08 μM RSL3 ± 1 μM BAZ (**C** for fluorescence microscopy images, scale bar = 60 μm; the left panel of **E** for analytical flow cytometry). **D** and the right panel of **E** are the respective quantitative intensity values (n = 3). **F.** Time-dependent induction of lipid-ROS accumulation following 0.08 μM RSL3 treatment (the left panel for flow cytometry assay; the right panel for quantitative intensity values, n = 3). **G, H.** Cellular levels of lipid-ROS after 8-h treatment with 0.08 μM RSL3 ± 1 μM BAZ (the left penal of **G** for flow cytometry assay, and **H** for confocal microscopy data, scale bar = 20 μm). The right panel of **G** for respective quantitative intensity values (n = 3). **I.** Levels of mitochondrial ROS after 8-h treatment with 0.08 μM RSL3 ± 1 μM BAZ (fluorescence microscopy images, scale bar = 60 μm). All quantitative data are presented as mean ± S.D. (* or ^#^ *P* < 0.05; ** or ^##^ *P* < 0.01).

In summary, these results indicate that BAZ has a strong protective effect against RSL3-induced ferroptosis in HT22 cells through attenuation of the accumulation of cellular NO, ROS, lipid-ROS and mitochondrial ROS.

### BAZ binds directly to PDI

The above observations prompted us to identify the cellular target that mediates the cytoprotective effect of BAZ. Our recent studies have shown that PDI is an important cellular protein that mediates chemically-induced, GSH depletion-associated oxidative cell death (20). Next we sought to determine whether PDI is a cellular protein that mediates the protective effect of BAZ against chemically-induced ferroptosis in HT22 cells. As summarized below, a number of experiments were performed.

We first determined the binding affinity of BAZ with PDI using the surface plasmon resonance assay, and found that BAZ could bind to PDI with a very high binding affinity (*K*_d_ = 3.3 nM based on curve fitting and 3.6 nM based on Scatchard plot) (**Fig. 6A, 6B**).

**Figure 6.**
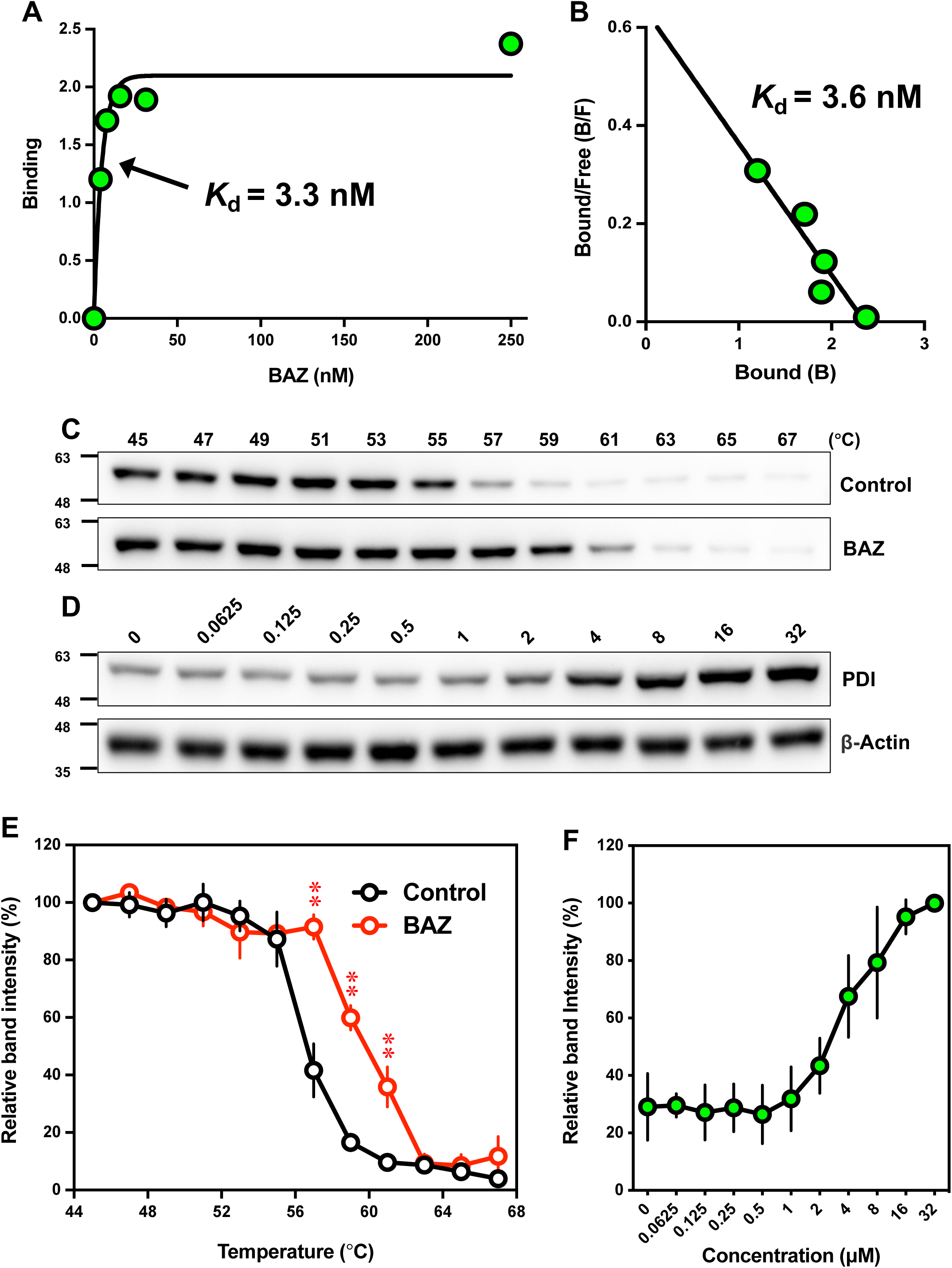
The binding of BAZ to PDI protein in live HT22 cells in culture. **A, B.** Surface plasmon resonance analysis of the binding affinity of BAZ for the purified PDI protein (**A** for the concentration-dependent binding curve; **B** for the Scatchard plot analysis based on the data in **A**). **C.** CETSA analysis of the change of PDI protein stability in BAZ (10 μM)-treated intact cells in response to increasing temperatures (n = 3). **D.** ITDR_CETSA_ analysis of the change of PDI protein stability in intact cells at 59°C in response to increasing concentrations of BAZ. **E.** The intensities of the protein bands in (**C**) were quantified using the Image J software. For CETSA curves, band intensities are calculated relative to the intensities at the lowest temperature for the BAZ-exposed samples and control samples, respectively (n = 3). **F.** For the ITDR_CETSA_ experiments, the relative band intensities were normalized to the β-actin samples (n = 3). All quantitative data are presented as mean ± S.D. (** *P* < 0.01).

Next, we performed the cellular thermal shift assay (CETSA) to evaluate the binding affinity between BAZ and PDI in live HT22 cells. Based on Western blot analysis of PDI protein stability, a thermal shift associated with PDI protein was observed in BAZ-treated HT22 cells compared to the control cells (**Fig. 6C**). A change in the Tm_50_ values (the temperature at which 50% of the proteins are precipitated by thermal denaturation) was determined to reflect the direct binding interaction of PDI protein with BAZ in cultured live cells. PDI had a Tm_50_ value of 56.5°C in control cells (in the absence of BAZ), but the presence of BAZ increased its Tm_50_ to 60°C, resulting in an increase in the Tm_50_ value (*i.e.*, ΔTm_50_) by 3.5°C (**Fig. 6E**).

To provide further support for the above experimental observations, the isothermal dose-response CETSA (ITDR_CETSA_) was also performed. The isothermal stability of PDI treated with different doses of BAZ at 59°C showed that PDI protein in live HT22 cells was stabilized by the presence of BAZ in a concentration-dependent manner (**Fig. 9D**). The CETSA curve nearly reached the plateau when 32 μM concentration of BAZ was present (**Fig. 9F**). Together, these experimental results show that BAZ can bind to PDI protein in live HT22 cells.

In this study, computational analyses were used to help understand the experimental observations on the PDILBAZ binding interactions. Based on computational docking analysis, 7 representative predicted structures of PDI□BAZ complexes are shown in **Fig. 7A** and **Supplementary Fig. S1A–S6A**. BAZ interacts closely with PDI’s ***b’*** domain and forms a hydrogen bond with His256 (**Fig. 7B, Supplementary Fig. S1B–S6B**). When PDI’s His256 is mutated to Ala256, the hydrogen bond formed between BAZ and His256 disappears (**Fig. 7D**, **Supplementary Fig. S1D–S6D**). The surfaces of the binding pockets are shown in **Fig. 7C** and **7E** (**Supplementary Fig. S1C–S6C, S1E–S6E**). The difference in the binding pockets of the wild-type and mutant PDI proteins is mostly in the region surrounding His256 which is mutated from a basic amino acid to a nonpolar amino acid (Ala256). The predicted binding affinities based on the 7 representative structures are summarized in **Table 2**. The binding energy values are increased after the HisIZAla mutation, indicating that the stability of the interaction between PDI and BAZ is decreased by the mutation.

**Figure 7.**
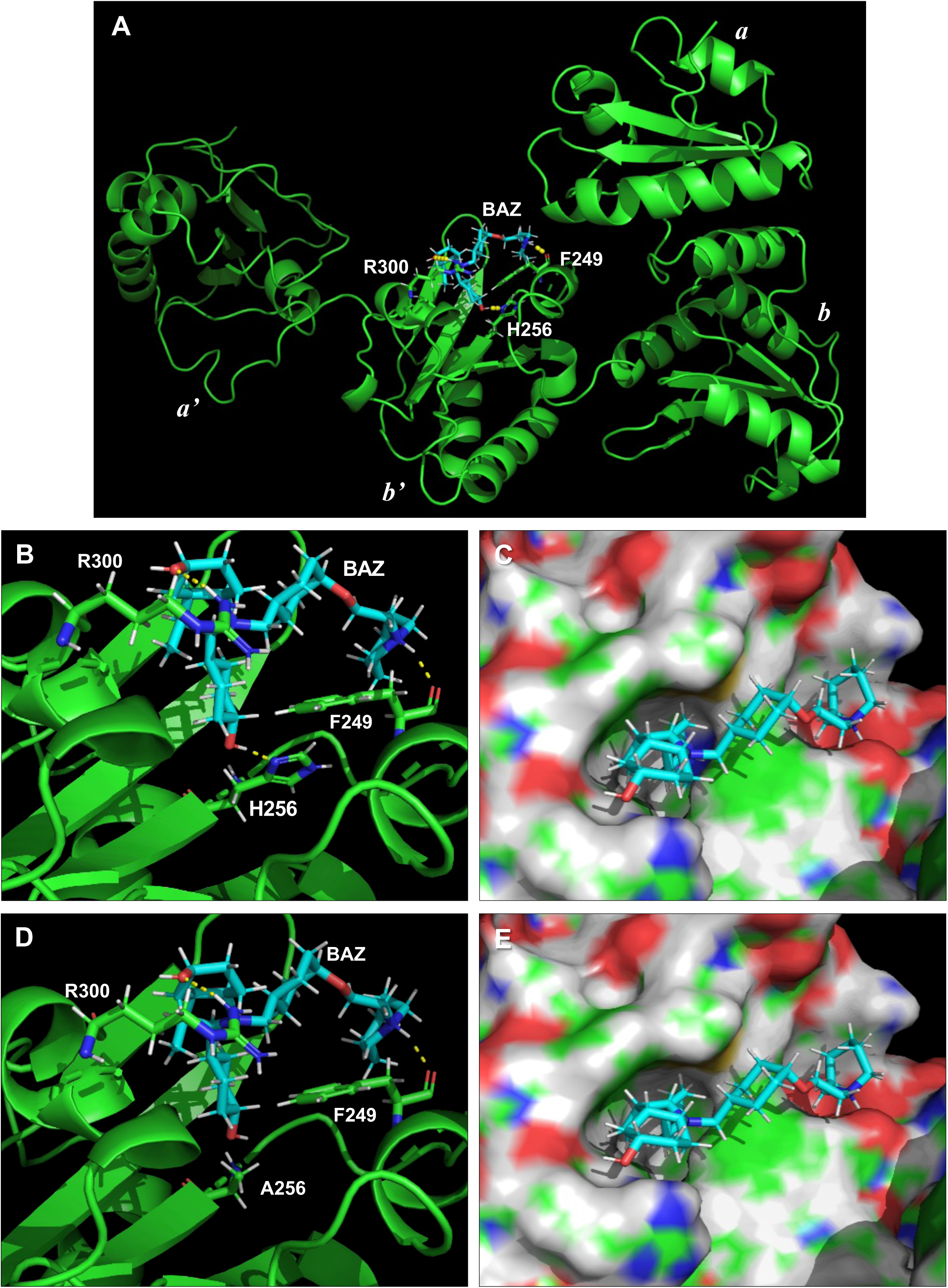
Predicted global structure of PDI □BAZ complex and the enlarged local structures of binding pockets. **A.** Predicted global structure of PDI□BAZ complex (representative docking pose #1). Three hydrogen bonds are formed between PDI and BAZ, and the involved residues (H256, F249 and R300) and BAZ are shown as sticks. PDI and BAZ are colored in green and cyan, respectively. **B** □**E.** Local structures of the binding pockets in the docking poses of the wild-type and the mutant PDI□BAZ complexes after MD simulation (**B, D**). Three hydrogen bonds are formed between BAZ and the wild-type PDI (**B**), but only two hydrogen bonds are formed between BAZ and the mutant PDI (**D**). In the local structure, the PDI and BAZ are colored in green and cyan, respectively. Panels **C** and **E** are the surface of the binding pockets of the wild-type and mutant PDI. The green regions are for carbon atoms, the blue regions for nitrogen atoms, the red regions for oxygen atoms, the white regions for hydrogen atoms, and the yellow regions for sulphur atoms, respectively.

**Table 2.** Predicted binding affinities based on the seven representative predicted structures of wild-type PDI□BAZ complexes or the Ala256 mutant PDI □BAZ complexes.

| PDI □ BAZ complex # | Number of hydrogen bonds between BAZ and PDI proteins | | Number of hydrogen bond between BAZ and His256 | | X-Score (Kcal/mol) | | PRODIGY-LIG (Kcal/mol) | | $\Delta_{\text{vinaRF20}}$ | |
| --- | --- | --- | --- | --- | --- | --- | --- | --- | --- | --- |
|  | PDI-His256 | PDI-Ala256 | PDI-His256 | PDI-Ala256 | PDI-His256 | PDI-Ala256 | PDI-His256 | PDI-Ala256 | PDI-His256 | PDI-Ala256 |
| 1 | 3 | 2 | 1 | 0 | -9.44 | -9.22 | -8.93 | -8.69 | 6.51 | 6.28 |
| 2 | 2 | 1 | 1 | 0 | -9.57 | -9.41 | -9.11 | -8.84 | 5.74 | 5.52 |
| 3 | 2 | 1 | 1 | 0 | -9.34 | -9.15 | -9.24 | -9.04 | 5.54 | 5.35 |
| 4 | 2 | 1 | 1 | 0 | -9.63 | -9.38 | -8.91 | -8.66 | 6.12 | 5.85 |
| 5 | 3 | 2 | 1 | 0 | -9.49 | -9.28 | -8.96 | -8.72 | 5.85 | 5.63 |
| 6 | 1 | 0 | 1 | 0 | -9.42 | -9.25 | -9.01 | -8.75 | 5.82 | 5.6 |
| 7 | 1 | 0 | 1 | 0 | -9.26 | -9.14 | -8.96 | -8.75 | 6.04 | 5.84 |

Next, MD simulation was performed to determine how tightly BAZ binds to the wild-type and mutant PDI proteins. In the case of wild-type PDI, it is observed that BAZ still remains bound to PDI in two of the three MD simulations (**Fig. 8A** and **8B**), but BAZ dissociates from PDI in one of the MD simulations (**Fig. 8C**). In comparison, for the mutant PDI-Ala256, BAZ dissociates from PDI in all three MD simulation trajectories (**Fig. 8G**, **8H** and **8I**).

**Figure 8.**
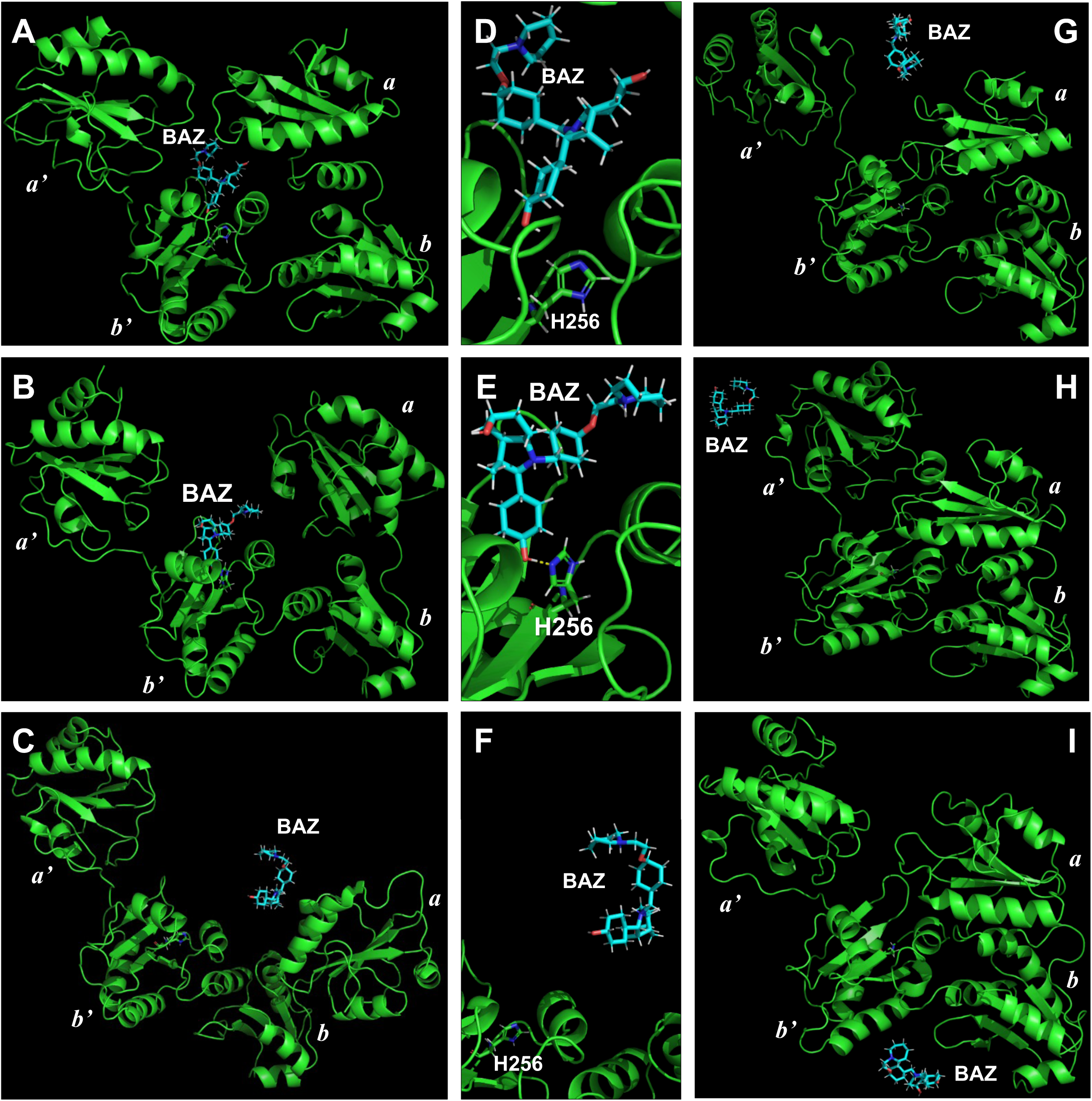
Conformations of the wild-type and mutant PDI □BAZ complexes after MD simulation. **A**□**C.** The last conformations the wild-type PDI□BAZ complexes selected along the first (**A**), second (**B**) and third (**C**) MD trajectories. The BAZ is still bound to PDI in the first (**A**) and second (**B**) cases, but it is dissociated from PDI in the third case (**C**). **D**□**F.** The local structures of the binding pockets in the last conformations of the wild-type PDI□BAZ complexes selected along the first (**D**), second (**E**) and third (**F**) MD trajectories. No hydrogen bond exists between BAZ and His256 in the first (**D**) and third (**F**) cases, and one hydrogen bond still exists between BAZ and His256 in the second case (**E**). **G**□**I.** The last conformation of the mutant PDI□BAZ complex along the first (**G**), second (**H**) and third (**I**) MD trajectories. Note that BAZ is dissociated from the mutant PDI-Ala256 in all three cases.

It is of interest to note that after MD simulation, the conformation of the wild-type PDI is changed from its initial open state (as shown in **Fig. 7A**) to a closed state (**Fig. 8A**) or partially-closed state (**Fig. 8B**). Similar observations are also made with the mutant PDI-Ala256, which is changed from its initial open state to the closed-state (**Fig. 8H**) to the partially-closed states (**Fig. 8G** and **8I**).

After MD simulation, the aromatic ring of BAZ remains buried inside the binding pocket of the wild-type PDI, whereas the other part of the BAZ molecule is rotated (**Fig. 7B**, **7C**, **Fig. 8D, 8E**). The contact number between BAZ and PDI (atom pairs within a given cutoff of 5 Å) was shown in **Supplementary Fig. S7A**. The contact number in the wild-type PDI□BAZ complex (**Supplementary Fig. S7A**) along two of the three MD trajectories fluctuates around 800 during the simulation process, and the number of contacts in the mutant PDI□BAZ complex along two of the three MD trajectories nearly goes down to zero after 40-ns MD simulation. While the minimum distance between BAZ and PDI-Ala256 becomes very long in the mutant PDI□BAZ complex in all three MD simulation trajectories, the distance between BAZ and PDI-His256 remains very close in two of the MD simulation trajectories (**Supplementary Fig. S8A**).

At the end of the 100-ns MD simulation (three trajectories), the hydrogen bond between BAZ and the wild-type PDI-His256 still exists in one complex (trajectory-2; **Fig. 8E**), and this hydrogen bond exists for a cumulative duration of 43.1 ns (**Supplementary S8B**). In another BAZ□PDI-His256 complex (**Fig. 8D**), BAZ moves slightly away from His256 at the end of MD simulation (trajectory-1), and the hydrogen bond exists for a cumulative duration of 18.5 ns (**Supplementary S8B**). In the third complex shown in **Fig. 8F**, BAZ moves away from the wild-type PDI from the start, and no hydrogen bond is formed (trajectory-3; **Supplementary S8B**). In contrast, for the three BAZ□mutant PDI complexes, no hydrogen bond formation is found during the 100-ns MD simulation.

It is observed that during MD simulations, the binding energies of the representative conformations of the wild-type PDI□BAZ complexes are generally lower than those of the mutant PDI□BAZ complexes, especially after 30-ns MD simulation (Supplementary Fig. S7B⍰S7D). The results are in line with the binding energy results for the seven representative PDI□BAZ complexes (docking structures) for both wild-type and mutant PDI (**Table 2**). Together, these results illustrate the relative importance of PDI’s His256 in its binding interaction with BAZ from both geometric and energy point of views.

### BAZ inhibits PDI’s catalytic activity

Results from our recent study (22) showed that the disulfide isomerase activity of PDI (which catalyzes the formation of disulfide bonds in target proteins in a cell) plays a role in mediating erastin-induced ferroptosis. Next, we sought to determine whether BAZ can directly inhibit PDI’s catalytic activity by using the *in vitro* insulin aggregation assay. Insulin has two chains cross-linked together by disulfide bonds, and PDI can break these disulfide bonds to facilitate the formation of aggregates, which has been commonly used to reflect PDI’s catalytic activity (37). We found that the PDI’s catalytic activity *in vitro* was suppressed by the presence of BAZ in a dose-dependent manner (**Fig. 9A**).

**Figure 9.**
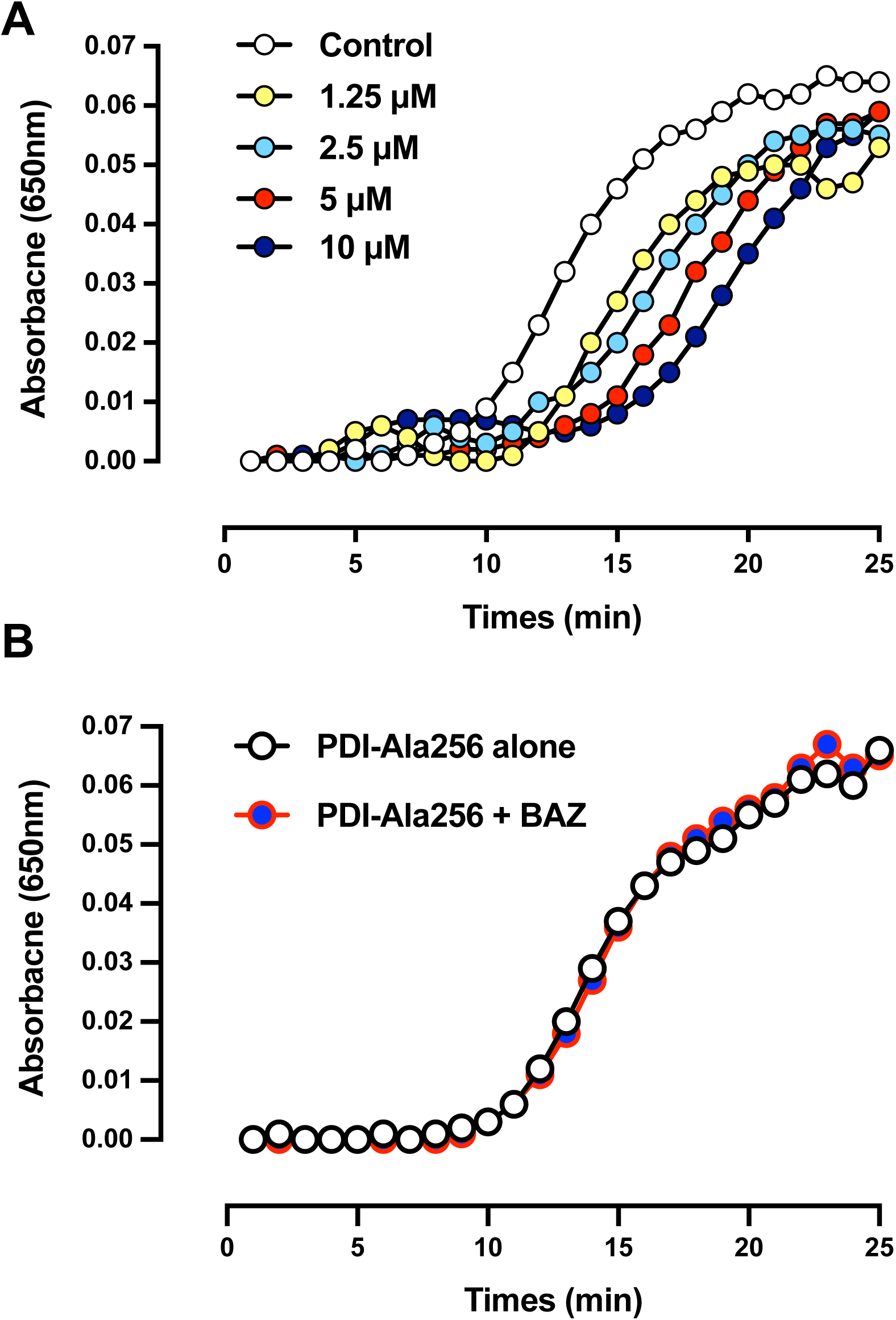
Effect of BAZ on the catalytic activity of the wild-type PDI and mutant PDI-Ala256. **A, B.** Inhibitory effect of BAZ on the catalytic activity of the wild-type PDI (**A**) and mutant PDI-Ala256 (**B**). PDI catalytic activity was assayed by measuring PDI-mediated insulin aggregation. Each experiment was repeated three times with similar results.

Our computational analysis with both wild-type PDI and mutant PDI-Ala256 has suggested that PDI’s His256 is important for the binding interaction between PDI and BAZ. To provide experimental support for the predictions based on computational analysis, next we chose to compare the ability of BAZ to inhibit the catalytic activity of the wild-type PDI and the mutant PDI-Ala256. As shown in **Fig. 9A** and **9B**, while the mutant PDI-Ala256 appeared to have a slightly-reduced catalytic velocity (*i.e.*, it took longer to catalyze the same reaction), its catalytic ability to complete the same maximal level of reaction (*i.e.*, in producing the same maximal level of insulin aggregation) was not changed. Notably, the presence of BAZ no longer showed a meaningful inhibition of PDI-Ala256’s catalytic activity in mediating the aggregation of insulin. This result indicates that BAZ cannot effectively bind to the mutant PDI-Ala256 to exert the same degree of inhibition of PDI-Ala256’s catalytic activity as it does with the wild-type PDI protein. Together, these results from the *in vitro* enzymatic assay demonstrate that BAZ can directly bind to the wild-type human PDI and inhibit its catalytic activity but it cannot do so with the mutant PDI-Ala256.

### PDI inhibition mediates BAZ’s protection against chemically-induced ferroptosis

Next, we sought to determine whether BAZ can effectively inhibit erastin/RSL3-induced NOS dimerization and accumulation of NO and ROS in HT22 cells through inhibition of PDI. Treatment of HT22 cells with BAZ abrogated erastin-induced increase in total NOS (nNOS and iNOS) proteins and NOS (nNOS and iNOS) dimer levels (**Fig. 10A, 10C**). In comparison, the PDI protein levels were not significantly altered by joint treatment with BAZ (**Fig. 10E**).

**Figure 10.**
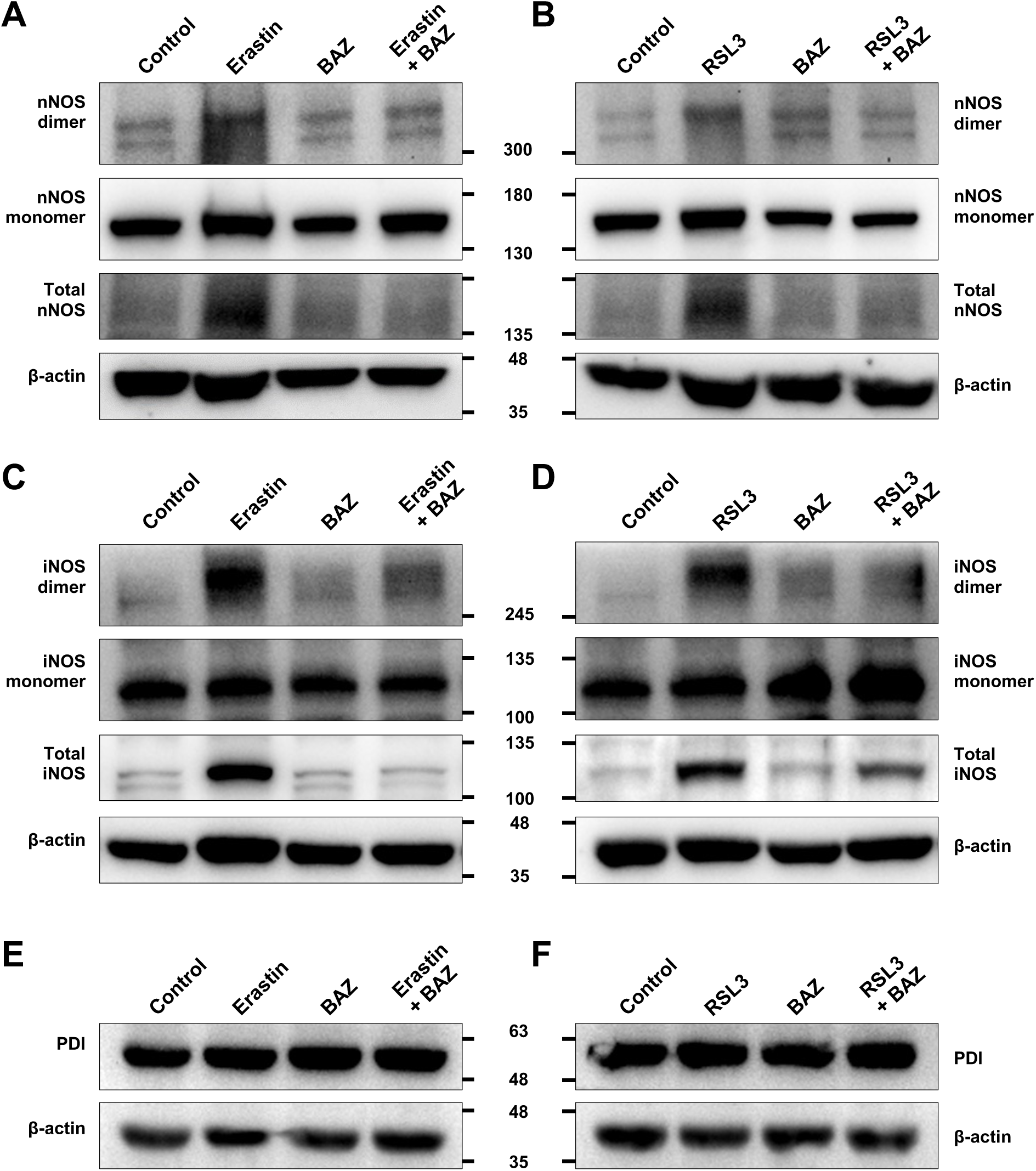
BAZ abrogates erastin- and RSL-induced increase in total nNOS and iNOS protein levels and their dimerization in HT22 cells. **A, B.** Levels of total cellular nNOS protein and its dimer and monomer forms following 8-h treatment with 0.8 μM erastin ± 1 μM BAZ (**A**) or 0.08 μM RSL3 ± 1 μM BAZ (**B**). The total cellular proteins (25 μg) were separated by 8% SDS-PAGE under reducing conditions or 6% SDS-PAGE under non-reducing conditions and then immunoblotted with specific antibodies for nNOS and β-actin. **C, D.** Levels of total cellular iNOS protein and its dimer and monomer forms following 8-h treatment with 0.8 μM erastin ± 1 μM BAZ (**C**) or 0.08 μM RSL3 ± 1 μM BAZ (**D**). The total cellular proteins (25 μg) were separated by 8% SDS-PAGE under reducing conditions or 6% SDS-PAGE under non-reducing conditions and then immunoblotted with specific antibodies for iNOS and β-actin. **E, F.** Levels of cellular PDI protein following 8-h treatment with 0.8 μM erastin ± 1 μM BAZ (**E**) or 0.08 μM RSL3 ± 1 μM BAZ (**F**). The total cellular proteins (25 μg) were separated by 8% SDS-PAGE under reducing conditions and immunoblotted with specific antibodies for PDI and β-actin.

Similarly, joint treatment of HT22 cells with BAZ similarly abrogated RSL3-induced increase in total NOS (nNOS and iNOS) protein and NOS (nNOS and iNOS) dimer levels (**Fig. 10B, 10D**), and the PDI protein levels were not significantly affected by the joint treatment (**Fig. 10F**).

### BAZ protects against kainic acid-induced memory deficit and hippocampal neuronal damage in mice

To determine whether BAZ has neuroprotective effects *in vivo*, we used the kainic acid-induced neuronal damage in the hippocampal regions of mice as an *in vivo* model, and the classical Y-maze test was used to evaluate the working memory in these animals. The basic assumption of the classical Y-maze test is that when an animal reaches the center of the maze and is faced with the choice of two new arms, it would tend to explore the arm which has not been explored right before (**Fig. 11B**). Based on this assumption, the theoretical alternation rate is thought to be 100% for a normal healthy mouse. However, we found that under real experimental conditions, the real alternation rate of the control mice (receiving saline vehicle injections only) was only 71.0 ± 2.6% (**Fig. 11C**), and this value is largely in agreement with some of the reported values (about 70%) in earlier studies (57,59,61). The alternation rate in kainic acid-injected mice was found to be around 50%. This value is as expected since the kainic acid-injected mice would have damaged hippocampus and impaired working memory and thus could not remember as well which arm it had already explored right before, and consequently, it would be more likely to make a random choice when faced with two options. The alternation rates of kainic acid-injected mice jointly treated with 3 and 5 mg/kg BAZ were 65.5 ± 1.8% (*P* < 0.01) and 64.1 ± 1.7% (*P* < 0.01), respectively, which were significantly higher than the corresponding rates in animals injected with kainic acid alone. However, in animals treated with the lowest dose (1.5 mg/kg) of BAZ, the animals had a slightly-improved alternation rate compared with kainic acid-injected animals (51.0 ± 3.2% *vs* 55.5 ± 2.4%; *P* = 0.026%) (**Fig. 11C**).

**Figure 11.**
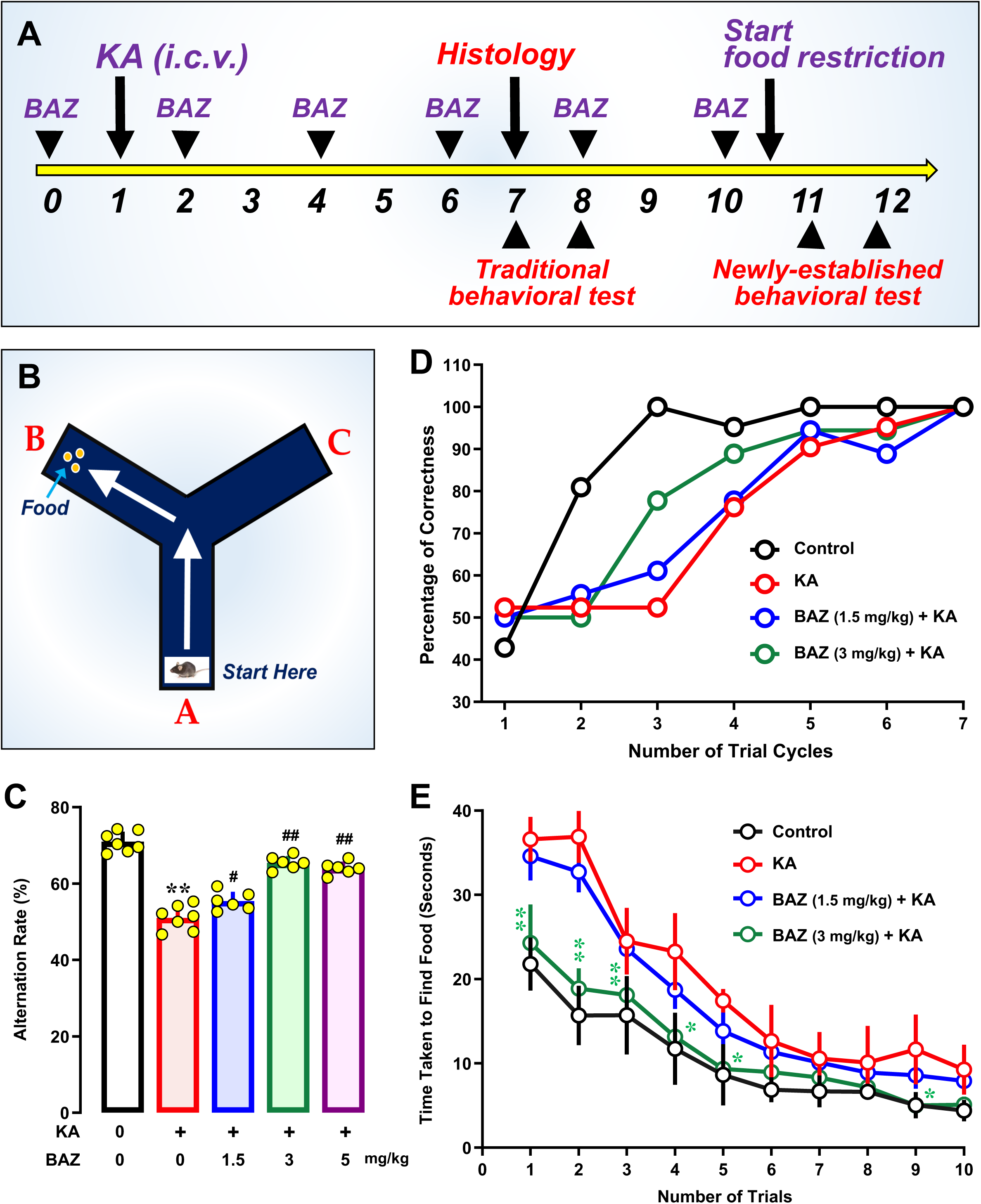
BAZ attenuates kainic acid-induced neurodegeneration in mouse hippocampus *in vivo*. **A, B.** The time scheme of the animal experiments (**A**) and the design of the Y-maze used in this study (**B**). **C.** Working memory of the animals in different treatment groups based on the alteration rates in the Y-maze test, conducted at 6 days after kainic acid (KA) treatment. **D.** Learning ability of the animals in different treatment groups assessed according to the correct rates (% of maximum) in finding the food and start eating it in different trial cycles. Note that the data from every three consecutive trials (except the last two trials) were used to calculate the mean value for one cycle. **E.** Learning ability of the animals in different treatment groups assessed according to the time (in seconds) taken to find the food and start eating it in the first ten trials. All quantitative data are presented as mean ± S.D. (n = 6□7; * or ^#^ *P* < 0.05; ** or ^##^ *P* < 0.01).

In the present study, a new Y-maze-based behavioral test was designed which enabled us to conduct additional assessment of the protective effect of BAZ on the learning and memory functions using the same kainic acid-injected mice. The procedures of this new behavioral test is described in detail in the methods section, and the data are summarized in **Fig. 11D**, **11E**. As shown in **Fig. 11B**, the food was placed in one arm of the Y-maze and the mouse was placed at the end of another arm (*i.e.*, the starting arm). Then the animal was allowed to freely explore the maze and would find the food and eat it. It was observed that nearly all the mice in the control group could correctly remember which arm contained food starting from the 7^th^ to 9^th^ trials and onward; in comparison, the kainic acid-injected animals took significantly more trials to correctly remember which arm contained food. Compared to the animals injected with kainic acid alone, those animals jointly treated with 3 mg/kg BAZ and kainic acid had a markedly improved memory, with close to 80% correctness between 7^th^IZ9^th^ trials and and >80% correctness thereafter (**Fig. 11D**).

For comparison, the overall time required for each animal to find the food and start eating it was also recorded in this behavioral test (**Fig. 11E**). It was observed that with increased number of trials, all mice in different experimental groups progressively improved their time, and eventually reached a similar plateau. This observation indicates that the animals in all treatment groups can gradually learn to perform this relatively simple task (*i.e.*, remembering the correct arm where the food is). However, in the first five trials, the differences among the four groups of animals were quite obvious, and there was a clear difference between the control group and kainic acid-injected group. In animals jointly treated with kainic acid + 1.5 or 3 mg/kg BAZ, the animals took significantly less time than animals injected with kainic acid alone, indicating that joint treatment with BAZ effectively improves the memory function of kainic acid-treated mice (**Fig. 11E**).

In this study, histological analysis of the representative hippocampal CA3 region of mice at 6 d post kainic acid injection was also analyzed to evaluate the neuroprotective effect of 3 mg/kg BAZ *in vivo*. Kainic acid-injected animals showed a drastic neuronal loss in the hippocampus, but joint treatment with 3 mg/kg BAZ for 7 d afforded almost complete neuroprotection in this brain region. The extent of neurodegeneration in the CA3 region was assessed using different methods (H/E, Fluoro-Jade B, and TUNEL staining) (**Fig. 12A–12C**). In H/E staining, kainic acid injection caused a strong loss of neuronal nuclei in the CA3 region, while this region in animals jointly treated with 3 mg/kg BAZ was strongly protected against kainic acid-induced neuronal death, almost making it not too different morphologically from the vehicle-treated control animals (**Fig. 12A**). This observation indicates that BAZ has a strong protective effect against kainic acid-induced oxidative neuronal loss *in vivo*. The observation made with the H/E-stained brain histology slides was also consistent with the slides stained with the Fluoro-Jade B and TUNEL assay kit. Fluoro-Jade B is an anionic fluorochrome capable of selectively staining degenerating neurons in brain slices (62). In the CA3 region in kainic acid-injected animals, it was found that a majority of neurons were strongly stained with Fluoro-Jade B (124.3 ± 17.5), whereas animals jointly treated with 3 mg/kg BAZ and kainic acid had a markedly reduced number of neurons in this region stained with Fluoro-Jade B (32.3 ± 8.7, *P* < 0.01) (**Fig. 12B**). Similarly, TUNEL assay selectively detects damaged cells that undergo extensive DNA degradation. Strong TUNEL signals in the kainic acid-injected group with mean fluorescence intensity of 69.2 ± 7.2 were detected, but the group jointly treated with BAZ had a markedly-reduced fluorescence intensity (35.9 ± 4.1, *P* < 0.01) (**Fig. 12C**). Together, these results demonstrate that joint treatment with BAZ offered a strong protection against kainic acid-induced neuronal death in a mouse model.

**Figure 12.**
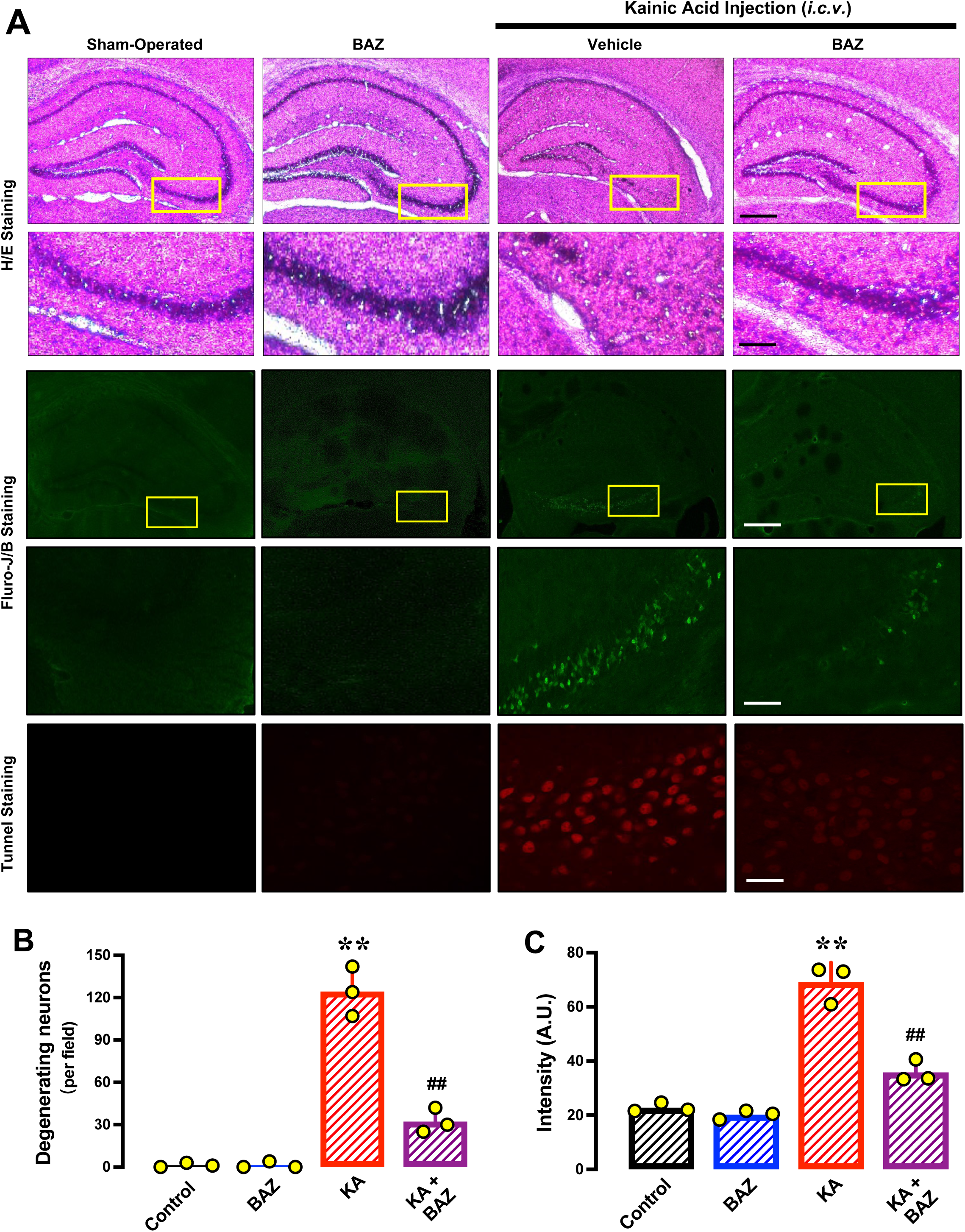
BAZ attenuates kainic acid (KA)-induced neurodegeneration in mouse hippocampus *in vivo*. **A.** Representative data for the histopathological analysis of the damaged hippocampal region. The upper two panels are the H/E staining of the hippocampal region (scale bar = 2 mm) and the enlarged CA3 region (scale bar = 200 μm). The middle two panels are the Fluoro-Jade B staining of the hippocampal region (scale bar = 2 mm) and the enlarged CA3 region (scale bar = 200 μm), respectively. The bottom panel is the TUNEL staining of apoptotic cells in the CA3 regions (scale bar = 60 μm). **B, C.** Quantitative data for the number of degenerating neurons (**B**) and TUNEL-positive cells (**C**) in control and kainic acid (KA)-treated mice (n = 3). All quantitative data are presented as mean ± S.D. (* or ^#^ *P* < 0.05; ** or ^##^ *P* < 0.01).

In summary, functional studies showed that mice treated with kainic acid alone have an impaired working memory compared to the control animals, but the impaired memory function is protected in a dose-dependent manner when the animals are jointly treated with BAZ. Histochemical analyses confirm that BAZ protects the hippocampal neurons of mice against kainic acid-induced neuronal injury.

## DISCUSSION

Recent studies have reported that some of the synthetic SERMs have a protective effect in neurons and glial cells against toxic insults (30-32). It has been suggested that the protective mechanism of SERMs may involve activation of ERs and G protein-coupled receptor for estrogens (GRP30) (33), which then triggers neuroprotective responses such as increasing the expression of antioxidants and activation of kinase-mediated survival signaling pathways. However, the exact mechanism underlying SERMs’ neuroprotective action is still unclear at present, which is the focus of our present study. We demonstrate that BAZ, a third generation SERM, exerts a strong protective effect against chemically-induced ferroptosis in hippocampal neurons both *in vitro* and *in vivo*.

Using the HT22 mouse hippocampal neuronal cells as an *in vitro* model, we show that treatment of these cells with erastin or RSL3 can readily induce cell death in a dose-dependent manner, and joint treatment of these cells with BAZ (at 62.5IZ1000 nM) can strongly protect the cells against erastin/RSL3-induced cell death in a concentration-dependent manner. The protective efficacy of BAZ is very high, *i.e.*, 100% protection against erastin-induced ferroptosis is observed when 500 nM BAZ is present, and similarly, 100% protection against RSL3-induced ferroptosis is observed when 125 nM BAZ is present. It is clear that the protective effect of BAZ against RSL3-induced cell death has a higher potency than its protection against erastin-induced ferroptosis.

The protective effect of BAZ against erastin- and RSL3-induced cell death is also evaluated in MDA-MB-231 human breast cancer cells, which do not express the estrogen receptors (63). It is evident that BAZ can exhibit a strong protective effect against erastin- and RSL3-induced ferroptosis in MDA-MB-231 cells, thus demonstrating that the protective effect of BAZ is independent of the estrogen receptor-mediated signaling pathways. In addition, we demonstrate that BAZ also has a strong protective effect against erastin- and RSL3-induced cell death in two other cell lines in culture (*i.e.*, the BRL-3A rat liver cell line and the H9C2 rat myocardium cells). While the apparent potency of BAZ in these cell lines are slightly different, its overall cytoprotective efficacy against chemically-induced ferroptosis is very similar to the observations made in HT22 cells.

The results of this study also demonstrate that BAZ can effectively protect against kainic acid-induced memory deficit and hippocampal neuronal damage in male mice. The male mice are selected for this study because the female mice normally produce and secret large amount of endogenous estrogens at cyclic fluctuating levels (depending their estrous cycle), and our earlier study has shown that the endogenous estrogens and some of their derivatives have a strong protective effect against neuronal cell death (60). Recently, we found that some of the estrogen derivatives can bind to PDI to exert their cytoprotective actions (unpublished data). As such, the use of male animals would help minimize this confounding factor. In addition, it is of note that several recent studies have shown that the kainic acid-neuronal cell death in the mouse brain closely resembles ferroptosis (64-66). It was reported that the brain tissue of kainic acid-treated mice has decreased levels of GSH, which is accompanied by increased levels of MDA, intracellular superoxide and lipid peroxides, increased PTGS2 expression and decreased GPX expression (64-66). All these changes are indicative of the induction of ferroptotic neuronal death.

Using the classical Y-maze test which is based on measuring the alternation rates of kainic acid-injected mice, it is observed that joint treatment of mice with 1.5, 3 and 5 mg/kg BAZ significantly improves the memory function. In this study, we also design a new Y-maze-based behavioral test to further assess the protective effect of BAZ on kainic acid-impaired learning and memory functions. It is shown that joint treatment of the animals with kainic acid + 3 or 5 mg/kg BAZ significantly improves the memory and learning performance. Histochemical analysis of the representative hippocampal CA3 region of mice 6 d post kainic acid injection shows a drastic neuronal loss, but joint treatment with BAZ (at 3 mg/kg) for 7 d affords almost complete neuroprotection in this brain region. The effect of BAZ in improving mouse learning and memory functions observed in this study is in agreement with earlier studies reporting a similar neuroprotective effect in other animal models (32).

To determine the mechanism of BAZ’s neuroprotective action, we hypothesize that BAZ may prevent erastin-induced cell death by targeting PDI, which subsequently inhibits PDI’s catalytic activity, reduces PDI-mediated conversion of nNOS monomers to its catalytically-active dimers, and ultimately, prevents chemically-induced ferroptotic cell death (as depicted in **Fig. 13**). As summarized below, experimental evidence is presented in this study to show that BAZ can bind directly to cellular PDI to inhibit its catalytic activity, then initiating a cascade of downstream changes culminating in ferroptotic cell death:

**Figure 13.**
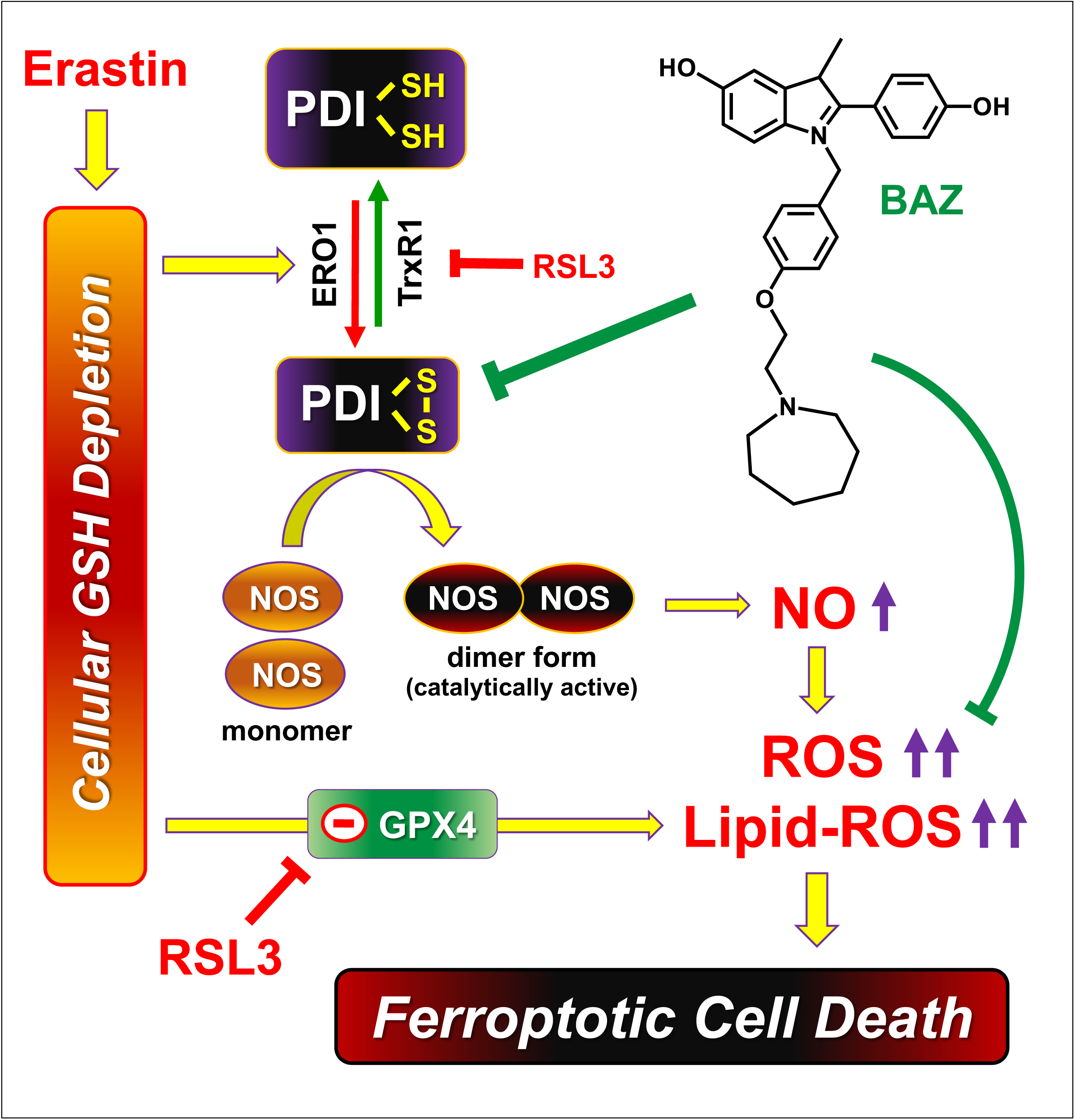
Schematic depiction explaining the mechanism by which PDI mediates the neuroprotective action of BAZ against chemically-induced ferroptosis. For detailed explanation, please refer to the Discussion section.

First, measurements based on the surface plasmon resonance technique show that BAZ can bind to purified PDI protein with a high binding affinity (apparent *K*_d_ of 3.3IZ3.6 nM). In addition, the cellular thermal shift assay (CETSA) show that BAZ can bind to PDI in live HT22 cells by causing an increase in the Tm_50_ value. In addition, the isothermal dose-response CETSA (ITDR_CETSA_) shows that PDI protein in live cells is stabilized by the presence of BAZ in a concentration-dependent manner. These experimental results jointly demonstrate that BAZ can strongly bind to PDI protein in live HT22 cells.

Second, computational docking analysis and MD simulations indicate that BAZ can bind tightly inside a rather deep binding pocket in the PDI protein, forming a hydrogen bond with His256. In addition, our docking results indicate that multiple Phe residues in the binding pocket may form hydrophobic interactions with the highly hydrophobic BAZ molecule, which contributes to the overall binding interaction between PDI and BAZ.

Here it is of note that during MD simulations, it is observed that the PDI protein undergoes an open-to-closed movement. In another recent MD simulation study, we also find that PDI can undergo open-to-closed or closed-to-open movements either in the absence or presence of small-molecule ligands (unpublished data). Since the residue-256 is not located in the hinge regions between different domains of PDI, it is suggested that the PDI-His256 residue (and its mutant PDI-Ala256 residue) likely would not affect the open-to-closed or closed-to-open movements of PDI.

Third, *in vitro* enzymatic assays show that BAZ can directly inhibit PDI’s catalytic activity *in vitro*. This direct inhibition of PDI’s catalytic activity is not observed with the mutant PDI-Ala256 protein. Interestingly, it is of note that while the catalytic velocity of the mutant PDI-Ala256 is slightly lower than the wild-type PDI, its maximal catalytic ability is still retained. Based on computational modeling analysis, the binding site of PDI for BAZ is located between its ***b*** and ***b’*** domains, whereas its catalytic site lies in the ***a*** and ***a’*** domains which contain the CXXC sequences. The reduced catalytic velocity of PDI-Ala256 is in full agreement with the notion that the His256 residue plays a role in the binding interaction with the protein substrates and also BAZ (an inhibitor). On the other hand, the mutant PDI-Ala256 still retains its maximal catalytic ability, which also fully agrees with the fact that the catalytic site (which lies between the ***a*** and ***a*’** domains) is not affected by the selective mutation and is also not affected by BAZ.

Together, these results showed that BAZ can directly bind to PDI, and the binding interaction is partly mediated through the formation of a hydrogen bond with His256. The binding interaction of BAZ with PDI is associated with an inhibition of PDI’s catalytic activity.

The finding that BAZ can bind tightly to PDI and effectively inhibit its catalytic activity *in vitro* is also supported by the biochemical changes observed in erastin-treated live HT22 cells. Specifically, it is observed that treatment of the cells with erastin alone results in PDI-mediated nNOS and iNOS activation (*i.e.*, formation of the dimer forms of nNOS and iNOS) along with cellular NO accumulation, and joint treatment of the cells with BAZ abrogates erastin-induced NOS dimerization and NO accumulation. In addition, BAZ also suppresses erastin-induced upregulation of NOS protein, which would also contribute to erastin-induced cytotoxicity. Together, these results demonstrate that BAZ can effectively protect against chemically-induced ferroptotic cell death partly by targeting PDI-mediated NOS dimerization and the subsequent accumulation of NO and lipid-ROS.

Based on the observations made in this study, it is evident that PDI is also involved, at least partially, in mediating RSL3-induced ferroptosis in HT22 neuronal cells, and inhibition of PDI’s function by BAZ contributes to its protection against RSL3-induced cell death. RSL3 is a prototypical ferroptosis inducer, and it has long been thought that its ferroptosis-inducing activity is primarily due to its inhibition of GPX4 (10). However, a recent study has reported that RSL3 can also strongly inhibit the enzymatic activity of thioredoxin reductase 1 (TrxR1) (67). Theoretically, inhibition of TrxR1 by RSL3 would shift the pool of cellular PDI proteins (a member of the thioredoxin superfamily) toward the catalytically-active oxidized state, thus favoring NOS dimer formation. Therefore, it was recently hypothesized by Hou et al. (65) that in addition to inhibition of GPX4, RSL3 may, through its ability to inhibit TrxR1 enzymatic activity (67), keep more PDI proteins in the oxidized state, thereby promoting RSL3-induced ferroptosis by facilitating NOS dimerization, followed by cellular NO, ROS and lipid-ROS accumulation, and ultimately ferroptotic cell death. A recent study has provided strong experimental evidence for this novel mechanistic explanation of RSL3-induced ferroptosis (65). This novel mechanism of RSL3-induced ferroptosis also provides a good explanation for the strong neuroprotective effect of BAZ against RSL3-induced ferroptosis, which is supported by experimental observations made in this study. For instance, we find that there is a time-dependent increase in total NOS protein levels in HT22 cells treated with 0.08 μM RSL3, and joint treatment of HT22 cells with BAZ abrogates RSL3-induced increase in total NOS protein and its dimer levels.

Lastly, it is of note that since BAZ exerts a markedly stronger protection against RSL3-induced ferroptotic cell death compared to erastin-induced cell death, it is possible that the direct antioxidant activity of BAZ against lipid-ROS may also contribute significantly to its cytoprotective effect. This possibility is in line with the fact that BAZ is a highly hydrophobic compound, and its antioxidant activity would be more readily manifested in protection against cellular lipid peroxides which are induced by treatment with RSL3. This suggestion is also consistent with an earlier study reporting that BAZ is a strong inhibitor of ferroptosis with a potent radical-trapping antioxidant activity (68). It is apparent that the results of our present study indicate that, in addition to its strong antioxidant activity, other mechanisms may also contribute importantly to BAZ’s strong cytoprotective action against chemically-induced ferroptosis.

As summarized in **Fig. 13**, the results of our study demonstrates that BAZ, a third generation SERM, exerts its strong protective effect against chemically-induced ferroptosis in hippocampal neurons both *in vitro* and *in vivo* through inhibition of PDI-mediated activation of NOS/NO in cells treated with erastin or RSL3, which subsequently results in reduction in cellular ROS and lipid-ROS levels, and ultimately ferroptotic cell death. In addition, the direct antioxidant activity of BAZ may also partially contribute to its protective effect against chemically-induced ferroptosis. These findings reveal a novel mechanism by which BAZ exerts its neuroprotective functions in an estrogen receptor-independent manner.

---

**Supplementary Figure S1.**
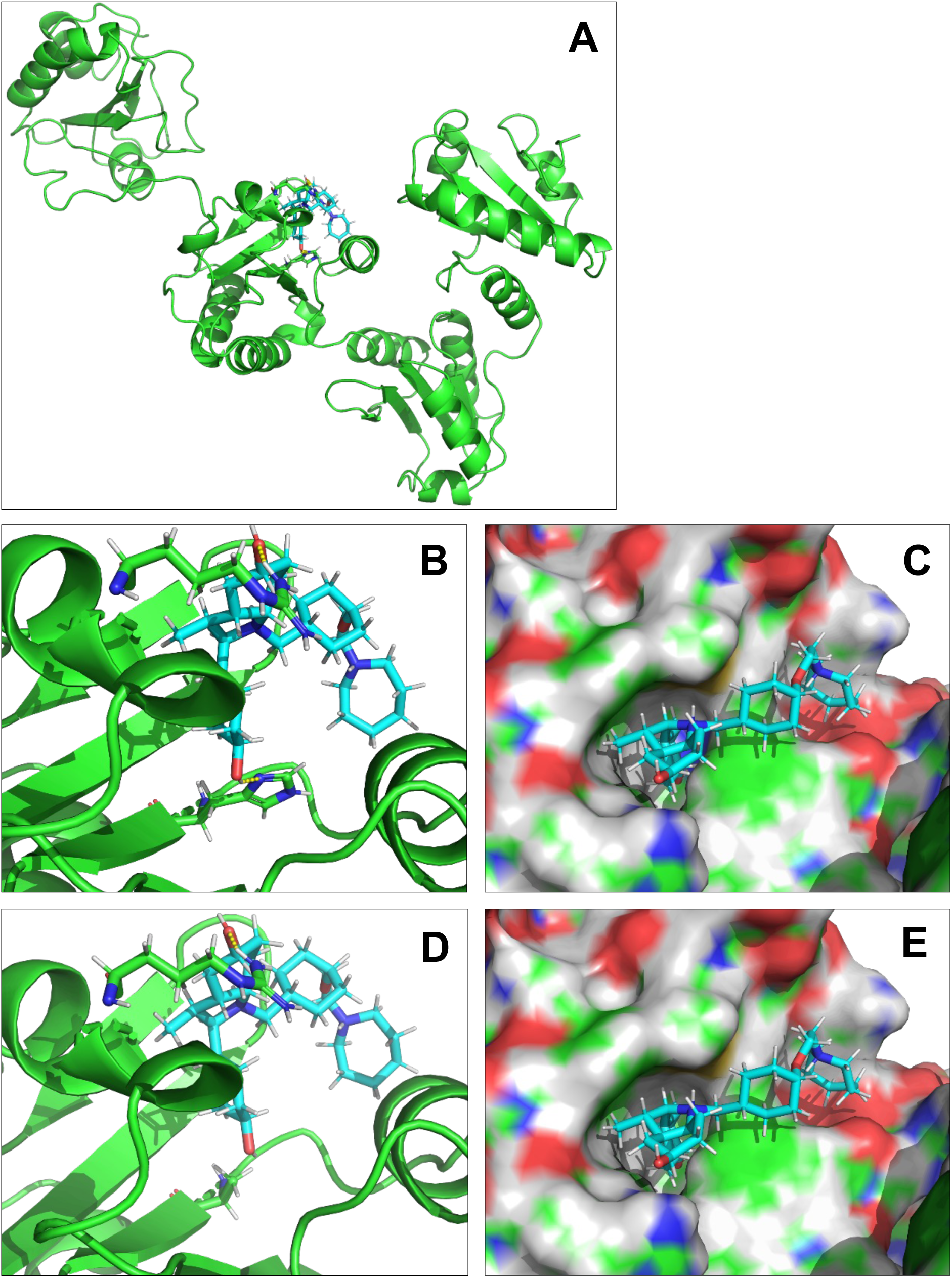

**Supplementary Figure S2.**
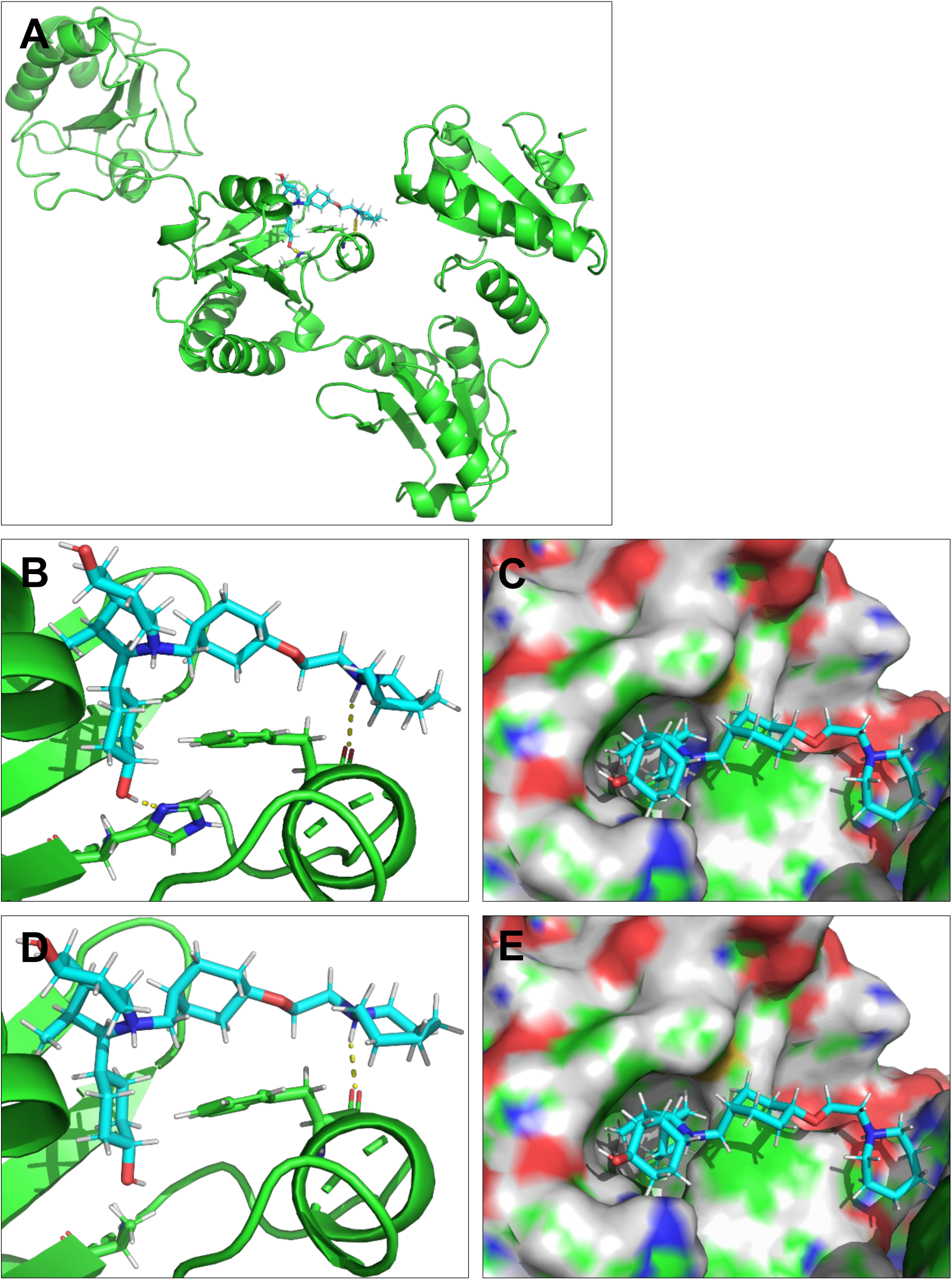

**Supplementary Figure S3.**
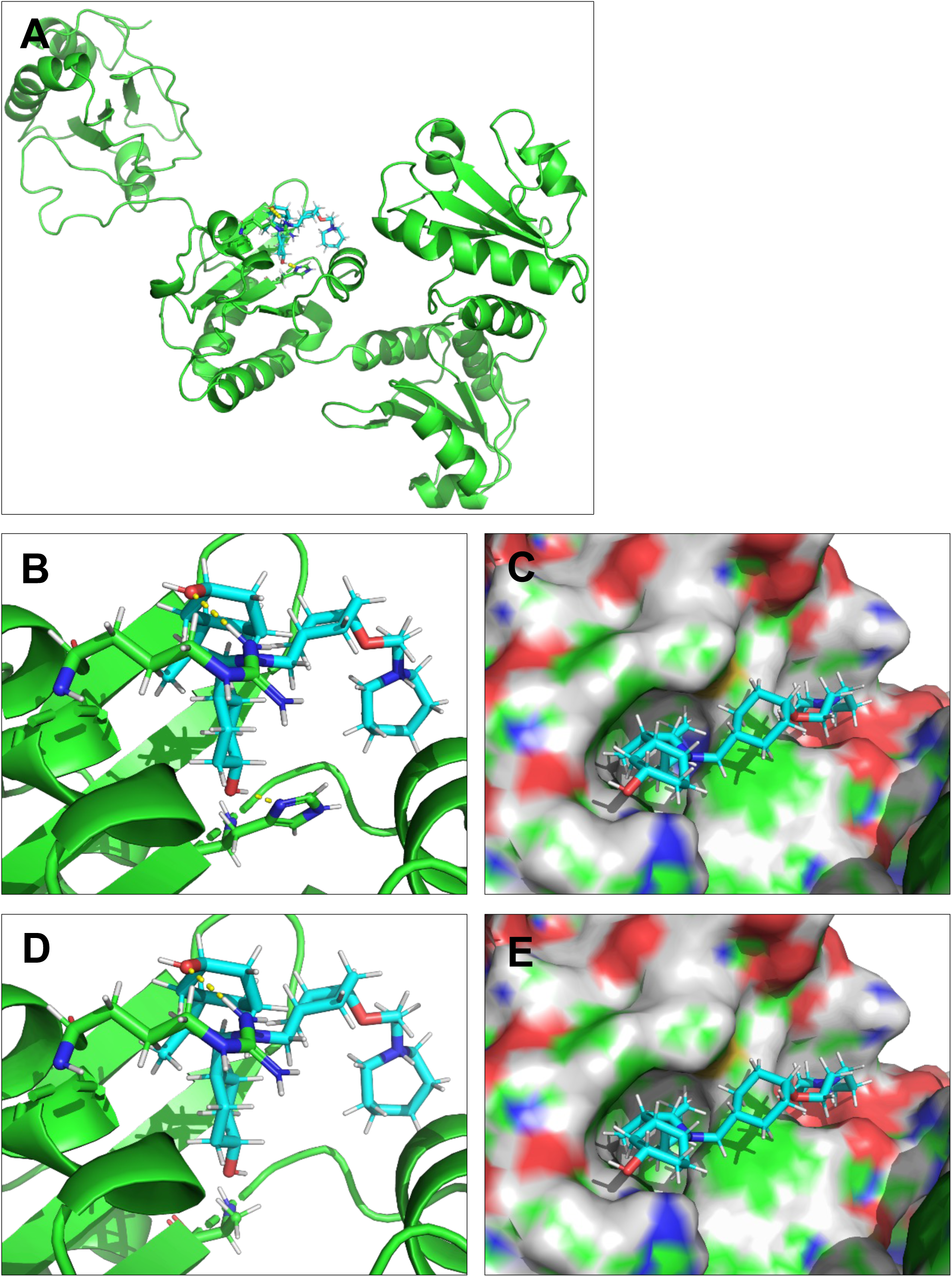

**Supplementary Figure S4.**
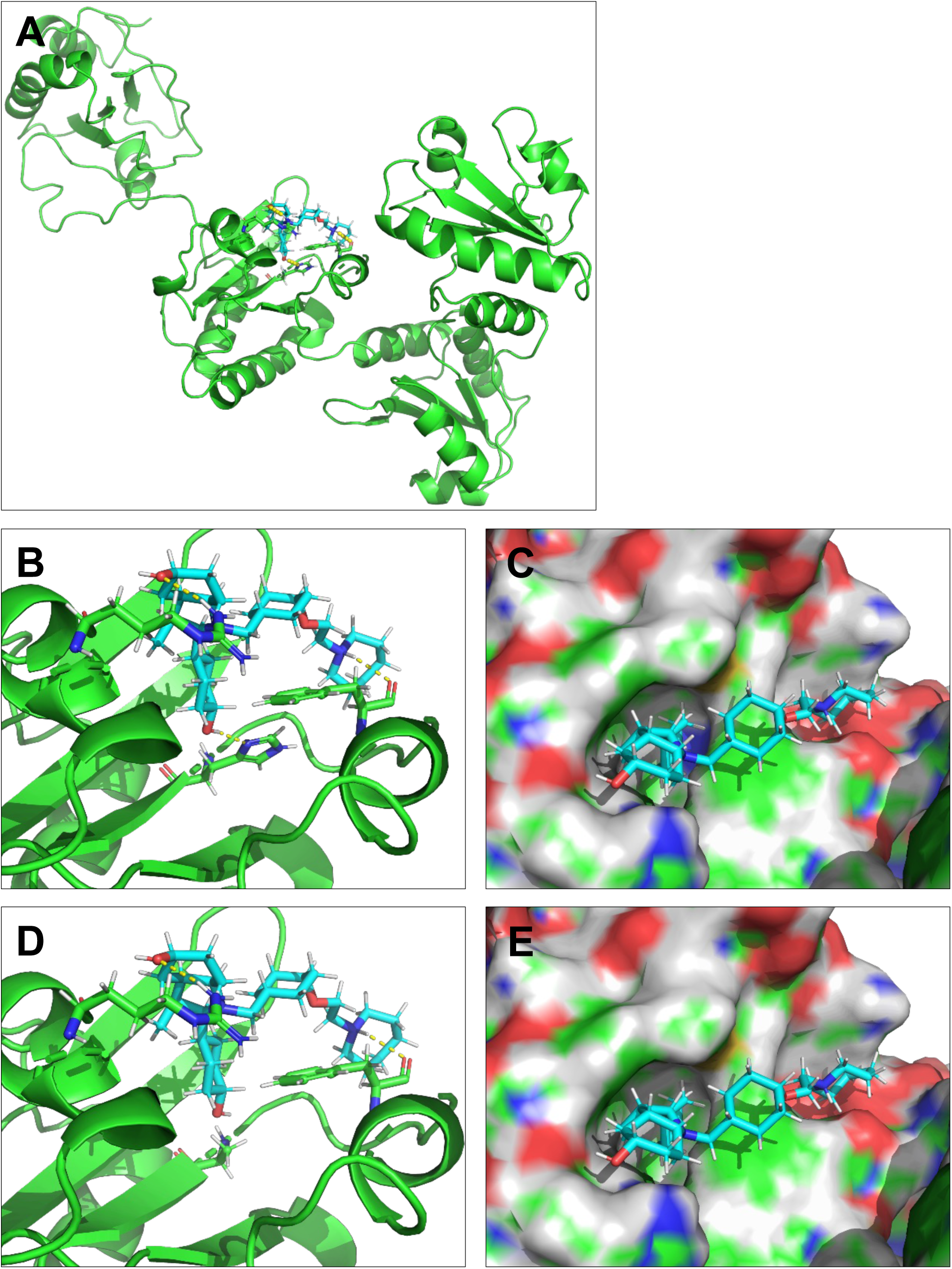

**Supplementary Figure S5.**
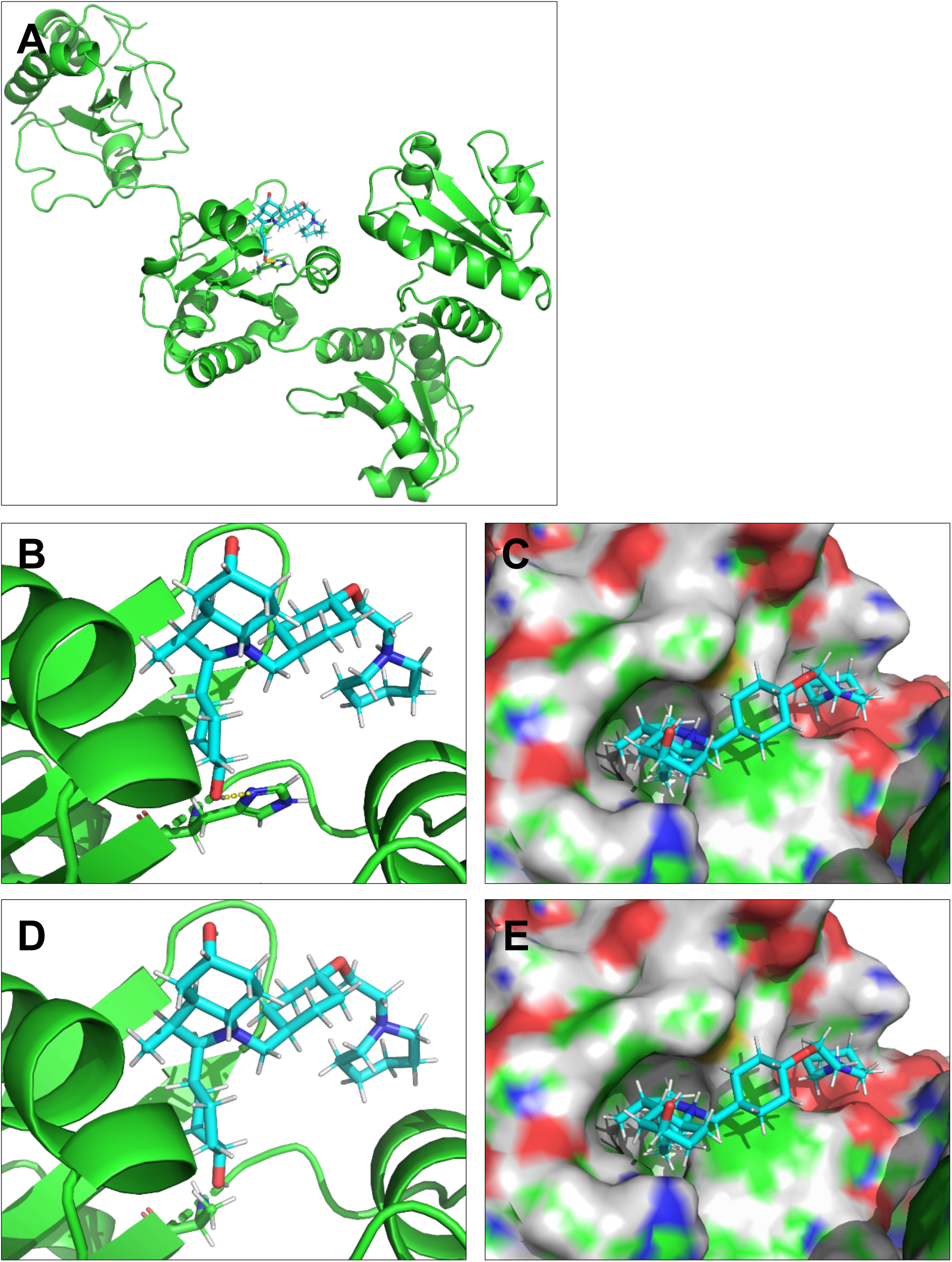

**Supplementary Figure S6.**
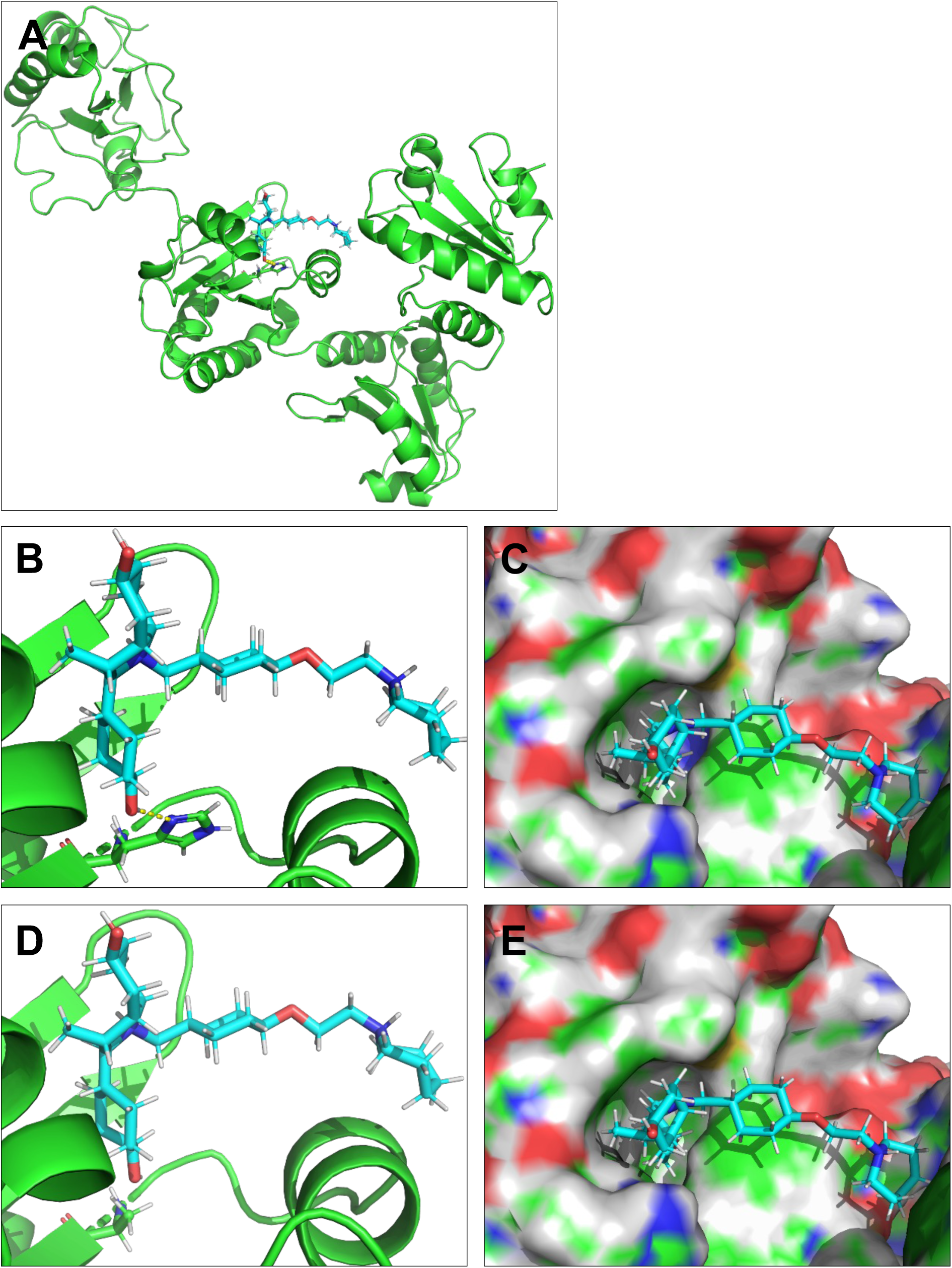

**Supplementary Figure S7.**
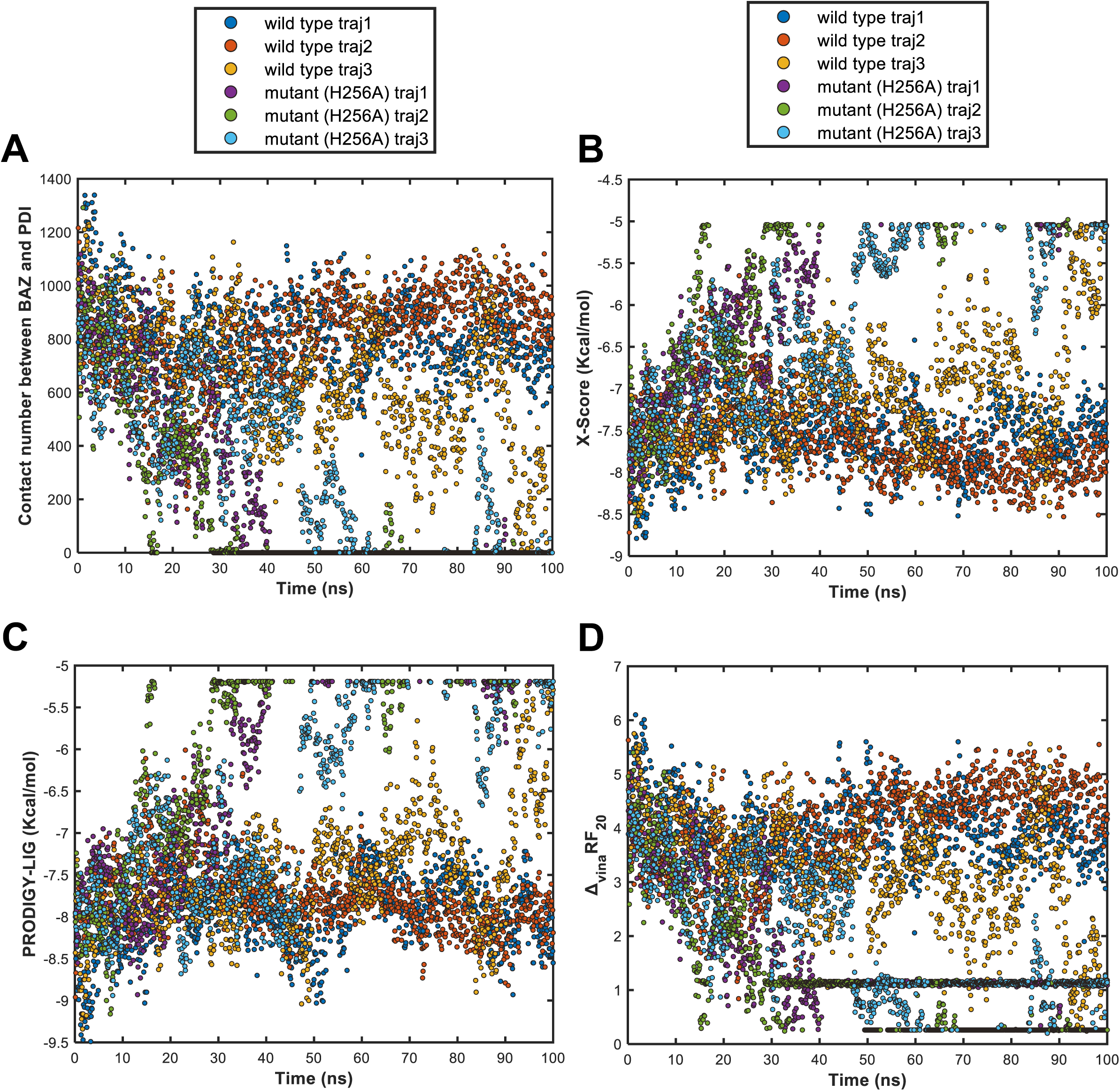

**Supplementary Figure S8.**
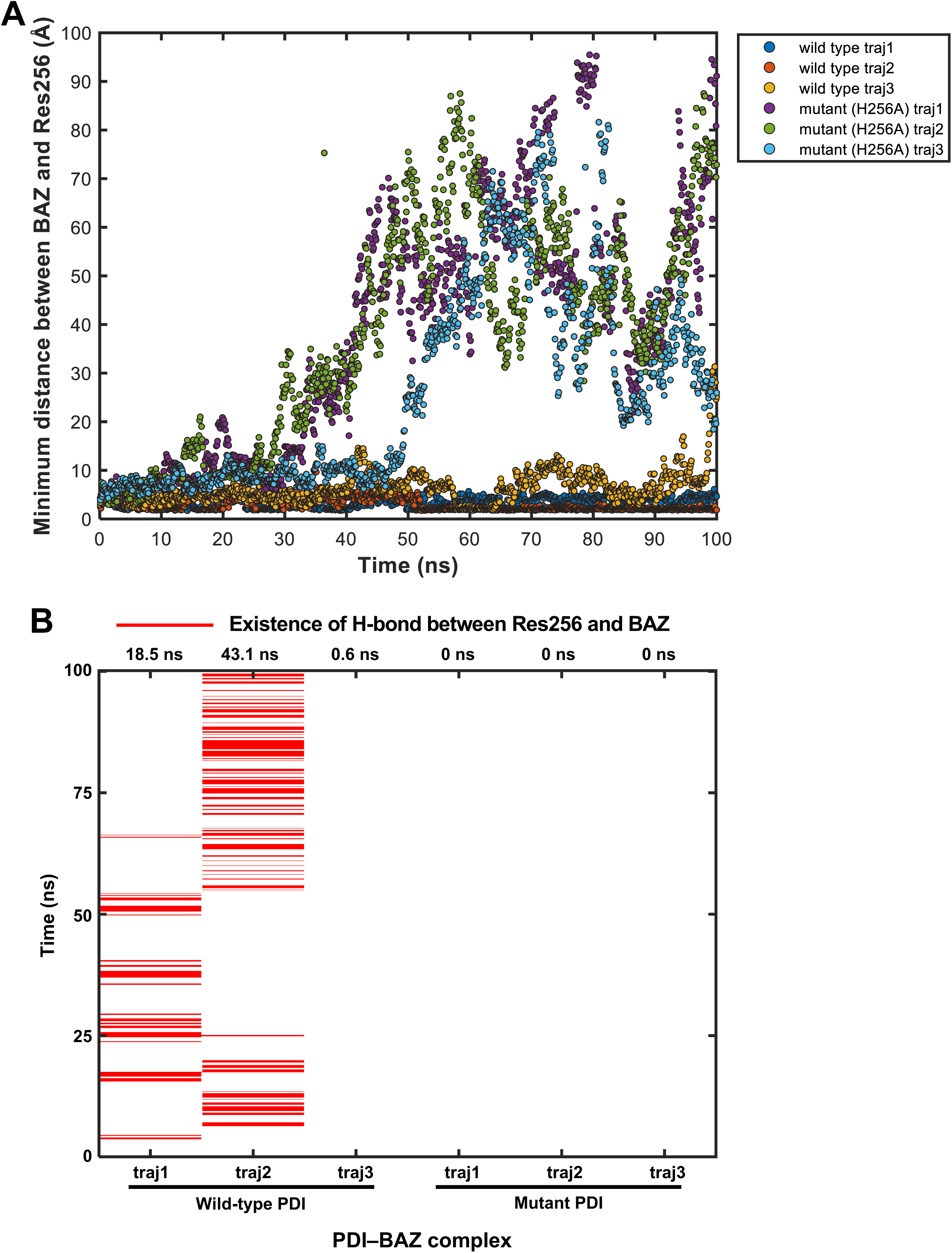

